# Endoplasmic reticulum exit sites are segregated for secretion based on cargo size

**DOI:** 10.1101/2023.12.07.570627

**Authors:** Sonashree Saxena, Ombretta Foresti, Aofei Liu, Stefania Androulaki, Maria Pena Rodriguez, Ishier Raote, Bianxiao Cui, Meir Aridor, Vivek Malhotra

## Abstract

TANGO1-family proteins (TANGO1, TANGO1S and cTAGE5) form stable complexes at the Endoplasmic Reticulum Exit Sites (ERES) and mediate export of bulky cargoes. The C-terminal proline rich domain (PRD) of these proteins binds Sec23A and affects COPII assembly at ERES. These PRD interactions were replaced with light-responsive domains to control the binding between TANGO1S-DPRD and Sec23A. TANGO1SΔPRD was dispersed in the ER membrane but relocated rapidly, yet reversibly, to pre-exiting ERES by binding to Sec23A upon light-activation. Prolonged binding of these two proteins concentrated ERES in the juxtanuclear region by a microtubule dependent process, blocked secretory cargo export and relocated ERGIC53 into the ER, but had limited impact on Golgi complex organization. Under these conditions, bulky collagen VII, and endogenous collagen I were collected at less than 47% of the stalled ERES, whereas small cargo molecules were halted uniformly across the ER, indicating that ERES differentially adapt to cargo size. We suggest these differences in cargo-accumulation at ERES permit cells to balance trafficking of cargoes of different sizes and optimize secretion.

## Introduction

A third of the cellular proteome enters the endoplasmic reticulum, and most of this pool is exported from specialized domains called ER exit sites (ERES).^1–3^ The export process is mediated by coat proteins (COPII) complex. ^4–6^ Sec12 located in the ER membrane, catalyzes the exchange of GDP for GTP on the cytoplasmic protein Sar1. Sar1-GTP is anchored to the ER membrane, and then recruits and assembles an inner and outer layer of COPII proteins.^7–10^ The inner layer, composed of Sec23/Sec24, selectively interacts with cargoes, cargo receptors and the outer coat layer (Sec13/Sec31). Recruitment of the outer layer stimulates GAP activity of Sec23 thus promoting GTP hydrolysis by Sar1.^11, 12^ During this process, COPII coated vesicles, of 60nm average diameter, filled with cargoes are produced for traffic to the next compartment of the secretory pathway.^13^

Clearly, these small vesicles cannot export bulky cargoes, like lipoprotein particles, collagens and other molecules, that compose the extracellular matrix.^14–18^ Considering that collagens constitute 17 % of mammalian dry weight ^19^, this is not a small problem.

Since cells export such a diverse collection of molecules, is there a division of labor amongst ERES to handle cargo load based on size and quantities? The identification of the proteins Transport and Golgi Organization 1 (TANGO1) gene-family members required for trafficking of bulky cargoes is helping to address this question.^20, 21^

Invertebrates possess only one Melanoma Inhibitory Activity (MIA) family gene, MIA3 encoding for TANGO1. In vertebrates the gene is duplicated to produce MIA2 (with two isoforms TALI and cTAGE5) and TANGO1 (with two isoforms TANGO1L and TANGO1S).^22, 23^ Complexes containing TANGO1, TANGO1S (a short isoform of TANGO1) and cTAGE5 bind Sec16 and Sec12, which in turn regulate the binding and hydrolysis of GTP by Sar1, thereby controlling COPII dynamics.^24^ TANGO1 consists of an ER-luminal domain for cargo binding, and a cytoplasmic part containing a proline-rich domain (PRD) for Sec23 binding.^21, 24–26^ TANGO1S and cTAGE5, like TANGO1, also bind Sec23 but they lack a luminal domain.^27^ TANGO1, together with its binding partners, TANGO1S and cTAGE5, forms a ring that compartmentalizes ERES within the ER membrane. TANGO1 also recruits ERGIC membranes to ERES, thereby creating a transient tunnel that can transport collagen from the ER lumen to presumably a tubular ERGIC emanating from the ER.^18, 27, 28^

To address the question of whether all ERES are the same with respect to the variety of cargoes exported from the ER, we have made use of optogenetics to control the binding of TANGO1S and Sec23A. We chose TANGO1S for our investigation because it retains the cytoplasmic sequence of TANGO1^29^, thus allowing us to elucidate effects of cytoplasmic interactions of TANGO1 on ER export. We observe that prolonged binding of these two proteins stalls cargo exit from the ER, concentrates ERES in the juxtanuclear region by a microtubule dependent process and causes relocation of ERGIC53 into the ER with limited perturbation to Golgi complex organization. In these conditions, exogenously expressed collagen VII and endogenous collagen I were collected at less than 47% of ERES, whereas small cargoes accumulated at most ERES thereby strongly indicating cargo selectivity at ERES.

## Results

### Manipulating TANGO1S and Sec23A binding by optogenetics

TANGO1S shares identical sequence with cytoplasmic domain of TANGO1 and is also required for collagen export from ER.^29^ TANGO1S binds Sec23A via its PRD domain and removal of this motif completely prevents their interaction. While TANGO1S only localize at ERES, TANGO1SΔPRD is dispersed throughout the ER membrane. Similar observations have also been reported for TANGO1ΔPRD (Full length TANGO1 without PRD).^21^ We employed optogenetics to control the binding of Sec23A to TANGO1SΔPRD. These optogenetic tools are based on an improved Light-Induced Dimer (iLID) system.^30^ A blue light sensitive Light-Oxygen-Voltage 2 (LOV2) domain is embedded with a peptide SsrA. The LOV2 domain undergoes a reversible conformational change in blue light and exposes the SsrA peptide. The now accessible SsrA peptide forms a dimer with its binding partner SspB. When blue light is removed, the domain undergoes a reversible conformational change, masking the SsrA peptide and causes a separation of the two peptides.^30–32^ As a blue light source, either a 488nm laser (for live imaging) or an in-lab designed blue LED set up (for fixed cells) was used. The N-terminus of Sec23A was tagged with EGFP-SspB while the C-terminus of TANGO1S and TANGO1SΔPRD were tagged with SsrA (also indicated as mCherry-iLID or mCh-iLID, **Figure 1A**). The modified TANGO1SΔPRD-mCh-iLID is localized uniformly throughout reticular and nuclear envelop membranes of the ER. The schematic in **Figure 1B** shows the expected effect of blue light on proteins generated by our procedures. During light-activation TANGO1SΔPRD-mCh-iLID binds to EGFP-SspB-Sec23A, and this dimerization is reversed in the absence of the blue light (**Figure 1B**).

**Figure 1:**
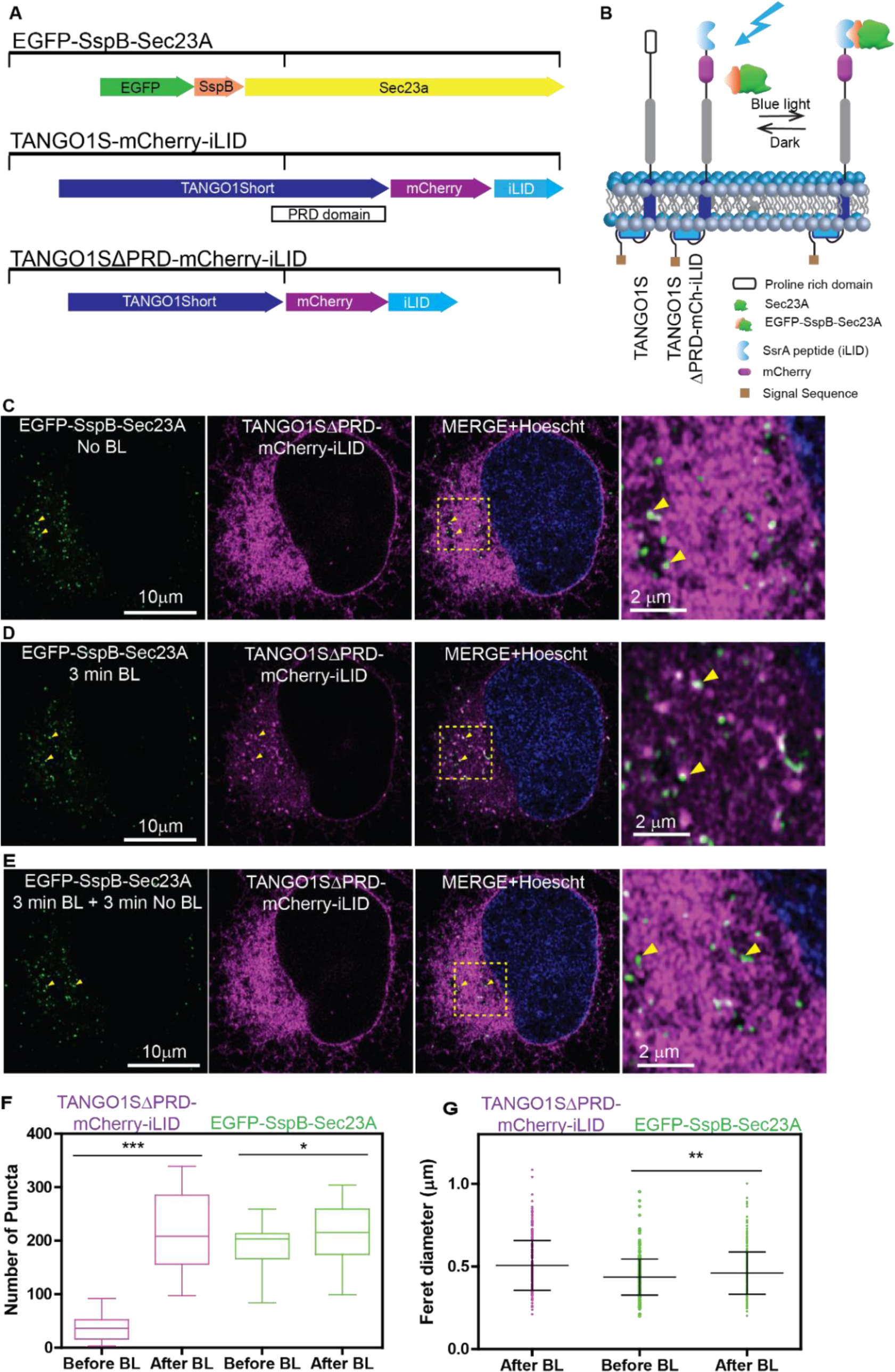
Manipulating TANGO1-Short and Sec23A binding by Optogenetics. **(A)** Schematics of optogenetic constructs of Sec23A, TANGO1Short (TANGO1S) and TANGO1SΔPRD. TANGO1S binds to Sec23A via its cytoplasmic proline rich domain (PRD). Deletion of PRD domain of TANGO1S (TANGO1SΔPRD) prevents its binding to Sec23A. The N-terminus of Sec23A is tagged with EGFP-SspB (SspB, is one half of an optogenetic pair). The C-terminus of TANGO1S is tagged with mCherry-iLID (iLID represents the second half of the optogenetic pair that contains SsrA). In the construct TANGO1SΔPRD, the PRD at the C-terminus of TANGO1S is replaced by mCherry-iLID. **(B)** Schematics of TANGO1S, TANGO1SΔPRD-mCh-iLID and EGFP-SspB-Sec23A as they align in the ER membrane. Blue light induces reversible dimerization of SsrA (iLID) and SspB domains and thereby binds TANGO1SΔPRD-mCh-iLID to EGFP-SspB-Sec23A. No blue light dissociates the complex of TANGO1SΔPRD-mCh-iLID and EGFP-SspB-Sec23A. **(C)** Snapshots of timelapse acquired on Airyscan (Supplementary material video 1A*) of live U2OS cells stably expressing TANGO1SΔPRD-mCh-iLID (magenta) and EGFP-SspB-Sec23A (green). In the absence of blue light (BL), TANGO1SΔPRD-mCh-iLID is dispersed throughout the ER membrane and EGFP-SspB-Sec23A is in discrete puncta. **(D)** Airyscan images of the same cell after 3 min exposure to blue light. TANGO1SΔPRD-mCh-iLID co-localizes within seconds with EGFP-SspB-Sec23A as seen in white puncta in overlay image and zoomed image (yellow arrowheads). **(E)** Airyscan images of the same cell further subjected to no blue light for 3 min. TANGO1SΔPRD-mCh-iLID is no longer visible at puncta and is observed throughout the ER membrane. **(F)** A box plot to compare the number of puncta formed by TANGO1SΔPRD-mCh-iLID (magenta) and EGFP-SspB-Sec23A, before and after exposing with blue light (BL) for 3 min. For TANGO1SΔPRD-mCh-iLID and EGFP-SspB-Sec23A statistically significant increase in number of puncta is seen after blue light exposure (p<0.0001, paired t-test) and (p<0.05, paired t-test) respectively. The horizontal line in each box corresponds to the mean, and error bars are shown. Data acquired from 15 cells in at least n=3 experiments. (Details in materials and methods). **(G)** An aligned dot plot showing the ferret diameters of puncta formed by TANGO1SΔPRD-mCh-iLID (magenta) after 3 min blue light exposure and EGFP-SspB-Sec23A (green) before and after 3 min blue light exposure. A statistically significant increase in the Feret sizes of EGFP-SspB-Sec23A was observed before and after BL exposure (p<0.01, paired t-test). Feret sizes of 300 distinct puncta were used from at least n=3 data sets. Details are provided in the materials and methods.

A stable U2OS cell line co-expressing TANGO1SΔPRD-mCh-iLID and EGFP-SspB-Sec23A was developed (Supplementary Figure 1A-C). In the absence of blue light (BL), EGFP-SspB-Sec23A is localized as discrete puncta and TANGO1SΔPRD-mCh-iLID is distributed throughout the ER. The zoomed overlay image on the right shows EGFP-SspB-Sec23 puncta, marked by yellow arrowheads (**Figure 1C**). When cells are exposed to blue light, the activation of SsrA (iLID) on the TANGO1SΔPRD-mCh-iLID enables it to bind to EGFP-SspB-Sec23A and localize to the ERES (**Figure 1D**). Live cell imaging revealed this reposition of TANGO1SΔPRD-mCh-iLID to EGFP-SspB-Sec23A puncta occurred within few seconds. The zoomed overlay image shows ERES marked by yellow arrowheads, which are now populated by both EGFP-SspB-Sec23A and TANGO1SΔPRD-mCh-iLID, observed as white puncta (**Figure 1D**, Supplementary Figure 1D). Within 3 min of turning off the blue light, the entire pool of TANGO1SΔPRD-mCh-iLID dissociates from EGFP-SspB-Sec23A and is dispersed throughout the ER. EGFP-SspB-Sec23A still remains localized to puncta (**Figure 1E**). Blue light irradiation did not have any effect on the stability of modified proteins in the cells (Supplementary Figure 1E). The average number of newly formed puncta of TANGO1SΔPRD-mCh-iLID (210 ± 70) was comparable to the average number of puncta of EGFP-SspB-Sec23A (212 ± 55) after exposure to blue light (**Figure 1F**). There was a small but significant increase in the number of puncta marked by EGFP-SspB-Sec23A before and after blue light, which suggested that TANGO1SΔPRD-mCh-iLID recruitment to puncta, was accompanied by enhanced assembly of EGFP-SspB-Sec23A (**Figure 1F**). The average Feret diameters of puncta of both the proteins were estimated to be around 0.5 μm, suggesting similar organization of the two (**Figure 1G**). To confirm that puncta marked by EGFP-SspB-Sec23A and TANGO1SΔPRD-mCh-iLID are ERES, we further investigated the composition of these sites.

### Real time recruitment of TANGO1SΔPRD to ERES by opto-genetics

Using the system described above, U2OS cells co-transfected with EGFP-SspB-Sec23A and TANGO1SΔPRD-mCh-iLID were exposed to blue light, fixed, and stained with antibodies against ERES markers (Sec16A and Sec31A), cTAGE5. The images revealed that the puncta formed by EGFP-SspB-Sec23A – TANGO1SΔPRD-mCh-iLID are enriched with these endogenous ERES markers (Supplementary Figure 2A-C) confirming that newly formed puncta are bona fide ERES.

To study the dynamic recruitment of ERES markers at these puncta, EGFP was removed from EGFP-SspB-Sec23A and a stable U2OS cell line co-expressing TANGO1SΔPRD-mCh-iLID and SspB-Sec23A was generated. Removal of EGFP did not affect the optogenetic tool functioning (Supplementary Figure 3A-D) as puncta formed by TANGO1SΔPRD-mCh-iLID were confirmed to be ERES marked by Sec16A and Sec31A (Supplementary Figure 3E).

The dynamics of ERES resident, Sec16, and outer layer of COPII complex, Sec13/Sec 31^13^, was studied during light-activation. Stable cells transfected with Sec16L-GFP were exposed to blue light. Sec16L-GFP was observed as discrete puncta at ERES (**Figure 2A**) while the ER-distributed TANGO1SΔPRD-mCh-iLID (**Figure 2A**) was recruited to Sec16L-GFP containing puncta after 3 min exposure to blue light (**Figure 2B**). A similar observation was made for stable cells transfected with Sec13-GFP, which was observed as discrete puncta as well (**Figure 2C**), and TANGO1SΔPRD-mCh-iLID populated these ERES with blue light (**Figure 2D**).

**Figure 2:**
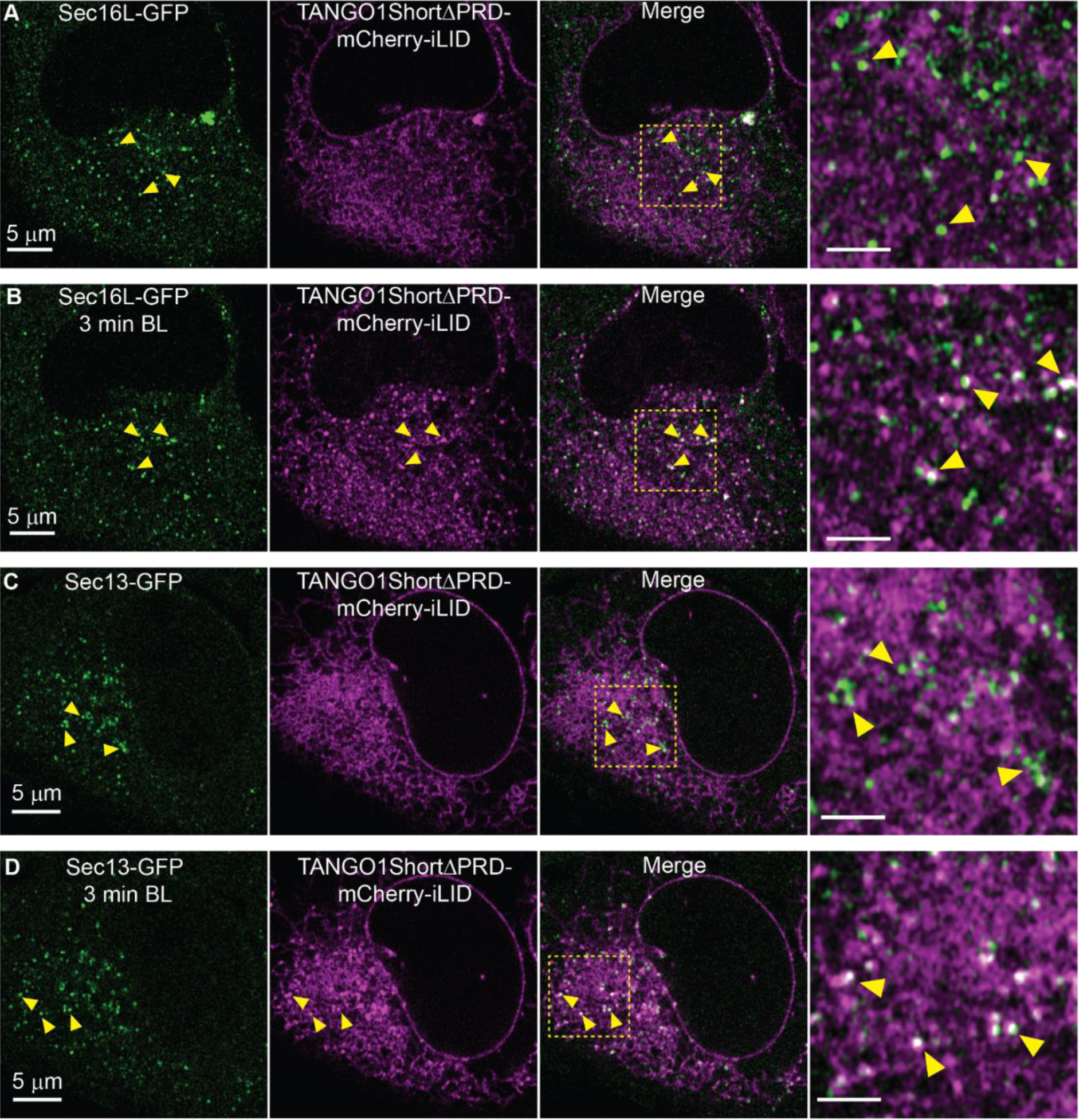
Real time recruitment of TANGO1SΔPRD to ERES by opto-genetics. Cells stably expressing TANGO1SΔPRD-mCh-iLID and SspB-Sec23A were transiently transfected with Sec16L-GFP or Sec13-GFP. Snapshots from time lapse (Supplementary material video 2A and 2B*) acquired on Airyscan are shown. **(A)** Sec16L-GFP (green) is observed in discrete puncta in the cells, whilst TANGO1SΔPRD-mCh-iLID (magenta) is dispersed in the ER membrane. Overlay and zoomed images (yellow arrowhead). Scale bar in zoom 2 μm. **(B)** After 3 minutes exposure to blue light, Sec16L-GFP (green) puncta are observed co-localized with TANGO1SΔPRD-mCh-iLID (magenta) as seen in white puncta in overlay image and zoomed image (yellow arrowheads). Scale bar in zoom 2 μm. **(C)** Sec13-GFP (green) is seen in discrete puncta, whilst TANGO1SΔPRD-mCh-iLID (magenta) is in the ER membrane. Overlay and zoomed images (yellow arrowhead). Scale bar in zoom 2 μm. **(D)** After 3 minutes exposure to blue light, Sec13-GFP (green) co-localized with TANGO1SΔPRD-mCh-iLID (magenta) as seen in white puncta in overlay image and zoomed image (yellow arrowheads). Scale bar in zoom 2 μm.

These experiments confirm that puncta formed by TANGO1SΔPRD-mCh-iLID were majorly a result of its accumulation at pre-existing ERES (marked by Sec16L and Sec13) while it remained bound to SspB-Sec23A. The average number of the ERES marked by Sec16L-GFP and Sec13-GFP were 300 ± 136 and 235 ± 38 respectively (Supplementary Figure 3F, G), and their average Feret diameters were ∼0.5μm (Supplementary Figure 3H, I) respectively, albeit enhanced recruitment of these component after blue light was not significant. These quantifications matched the features of puncta observed for TANGO1SΔPRD-mCh-iLID (**Figure 1F, G**). Overall, our data suggests that we achieved spatial and temporal control over the binding between TANGO1S and the COPII inner layer (Sec23) while both localize to ERES.

### Optogenetically formed ERES undergo fusion and fission and their formation requires Sar1 activity

ERES have been previously shown to undergo fusion and fission events.^33^ Similar events were also observed in our system. Upon exposure to blue light, the discrete puncta formed by accumulation of TANGO1SΔPRD-mCh-iLID and EGFP-SspB-Sec23A exhibited dynamic fission and fusion events (**Figure 3A-3C**). It has been reported that the fusion within two distinct ERES are identified by an increase in fluorescence intensity of Sec23A post fusion.^34^ In our observations of visible ERES coalescing events, the mean gray values of EGFP-SspB-Sec23A and TANGO1SΔPRD-mCh-iLID per ERES (post fusion) revealed a ∼1.5-fold increase in the fluorescence intensity, implicating an increase in number of each protein **(Figure 3D)**. Therefore, the persistent association of TANGO1SΔPRD-mCh-iLID and EGFP-SspB-Sec23A at ERES, driven by blue-light does not alter the dynamic nature of ERES. For this reason, our data suggest that the fusion events do not rely on the cycle of GTP binding and hydrolysis by the small GTPase Sar1 and rather are more akin to the physical behavior of protein-protein phase separation at ERES.^35, 36^

**Figure 3:**
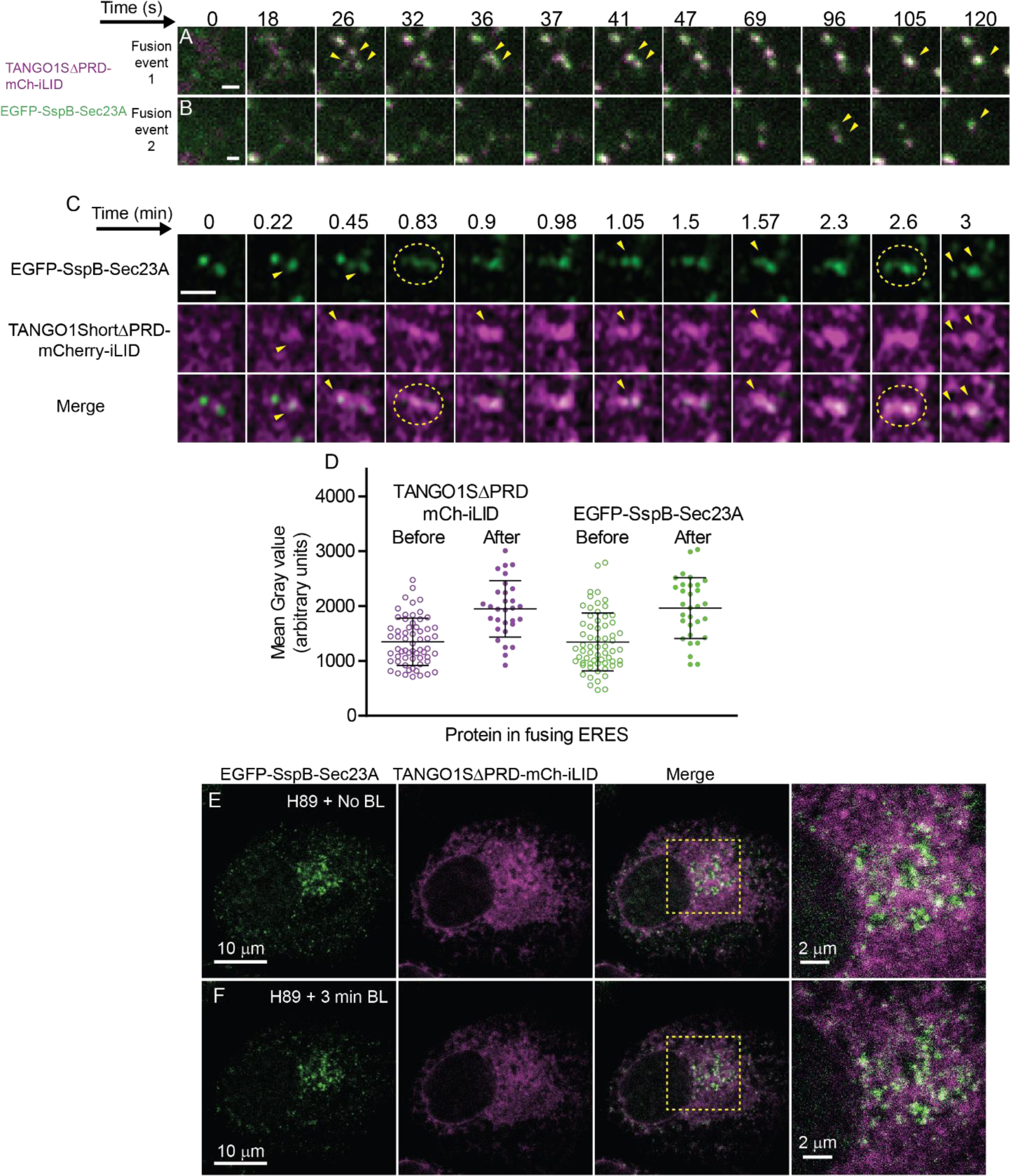
ERES undergo fusion and fission and their formation requires Sar1 activity. **(A, B)** Snapshots of a 2 min timelapse (Supplementary material video 3A*) of U2OS cell co-expressing TANGO1SΔPRD-mCh-iLID (magenta) and EGFP-SspB-Sec23A (green). Shown are two representative fusion events (yellow squares) observed during imaging of one cell in a spinning disk microscope. In fusion event 1 (**A**), three distinct puncta (marked by yellow arrow heads) are observed in close proximity at 26s. Within 10 seconds they form a single structure (time 36s). At 41s, the structure separates into two punctum and finally merges back into one punctum at time 105s and remains as such until the end of time lapse (120s). The fusion event 2 (**B**), shows similar kinetics where two punctate merges at 120s (shown by yellow arrowheads). Scale bar 1 μm. **(C)** Snapshots of a 3 min time lapse (Supplementary material video 3B*) of U2OS stably co-expressing EGFP-SspB-Sec23A (green) and TANGO1SΔPRD-mCh-iLID (magenta) acquired by Airyscan microscopy. Yellow dotted ovals show fusing ERESs, and yellow arrowheads show the puncta of interest. Three distinct ERES visible at 0.45 min in green channel are observed to form a merged continuous structure (0.83 min) marked by yellow dotted ovals. A similar structure is observed in magenta channel too, showing TANGO1SΔPRD-mCh-iLID forming a continuous structure as well. The ERES separates back into three distinct puncta (1.05 min) and these puncta again form a continuous structure at 2.6 min, which further breaks into discrete puncta at 3min. Scale bar 1 μm. **(D)** Plots of the mean intensity of TANGO1SΔPRD-mCh-iLID and EGFP-SspB-Sec23A accumulated at the discrete puncta before (hollow green and magenta circles) and after (filled green and magenta circles) fusion during blue light exposure for 2 min. The mean intensity increased ∼ 1.5 folds for each protein (30 fusion events quantified from Supplementary material video 3A*). **(E, F)** U2OS stably co-expressing TANGO1SΔPRD-mCh-iLID and EGFP-SspB-Sec23A were treated with 50μM H89 for 1 min prior to imaging. Cells were imaged before (**E**) and after (**F**) 3 min exposure to blue-light and the time lapse was acquired on confocal microscope (Supplementary material video 3C*). TANGO1SΔPRD-mCh-iLID (magenta) in the ER membrane and the EGFP-SspB-Sec23A (green) diffused in non-discrete puncta. Fewer ERES were observed with diminished intensity of EGFP-SspB-Sec23A, primarily located in the juxtanuclear region.

We also observed that ERES transiently associate into continuous structures (indicating fusion) that contain both TANGO1SΔPRD-mCh-iLID and EGFP-SspB-Sec23A (**Figure 3C**, yellow dotted oval).

The recruitment of Sec23/Sec24 to the ER membranes begins with insertion of Sar1-GTP into the outer leaflet of the ER membrane.^8^ H89 inhibits the recruitment of Sar1^7^ and its activation on ER membranes,^7, 37, 38^ directly affecting the formation of ERES (**Figure 3E**). Exposure to blue light in H89 treated cells, did not induce the formation of puncta of TANGO1SΔPRD-mCh-iLID and EGFP-SspB-Sec23A, the former remained dispersed in the ER membrane and was not collected in discrete puncta (**Figure 3F**). Therefore, the formation of puncta by light activation depends on the activation of Sar1, which initiates coat recruitment.

### ERES are mobilized to MTOC upon prolonged TANGO1SΔPRD and Sec23A binding

Prolonged expression of Sar1 mutants that are deficient in GTP hydrolysis leads to the clustering of ERES at the MTOC.^39^ Microtubule (MT) molecular motors dynein bind Sec23 using the p150glued-containing dynactin complex, therefore pulling ERES towards the MTOC.^40^ However, it is not known whether the clustering of ERES under prolonged over expression conditions is derived from stabilization of selective ERES at MTOC or if these are actively mobilized within the ER membrane by MT motors to that location. The highly dynamic nature of COPII coat at ERES negates prolonged COPII-MT interactions and long-range movement. However, ERES at the juxtanuclear region close to Golgi are more abundant in number, and hence many ERES are visible in that region (**Figure 4A**). Here we suggest that prolonged binding of TANGO1S to Sec23 upon blue light exposure potentially stabilizes dynein-ERES interactions. This lack of dynamics led to the accumulation of ERES in the juxtanuclear region, now visible as fused structures (**Figure 4B**). Treatment with nocodazole which depolymerizes microtubules, does not alter the binding of TANGO1SΔPRD-mCh-iLID to EGFP-SSPB-Sec23A, upon blue light exposure (**Figure 4C**). However, the ERES remain spread out and do not move to the juxtanuclear region, indicating their movement in cells is microtubule dependent (**Figure 4C, D**). The results suggest that COPII-MT motor binding leads to the mobilization of TANGO1S-containing ERES toward the MTOC. The physiological roles for an “in membrane” movement of ERES in sorting and traffic remains to be defined. The results further suggest that COPII dynamics is inhibited during light activation. We therefore studied the consequences of stabilizing the ERES on cargo sorting and traffic.

**Figure 4:**
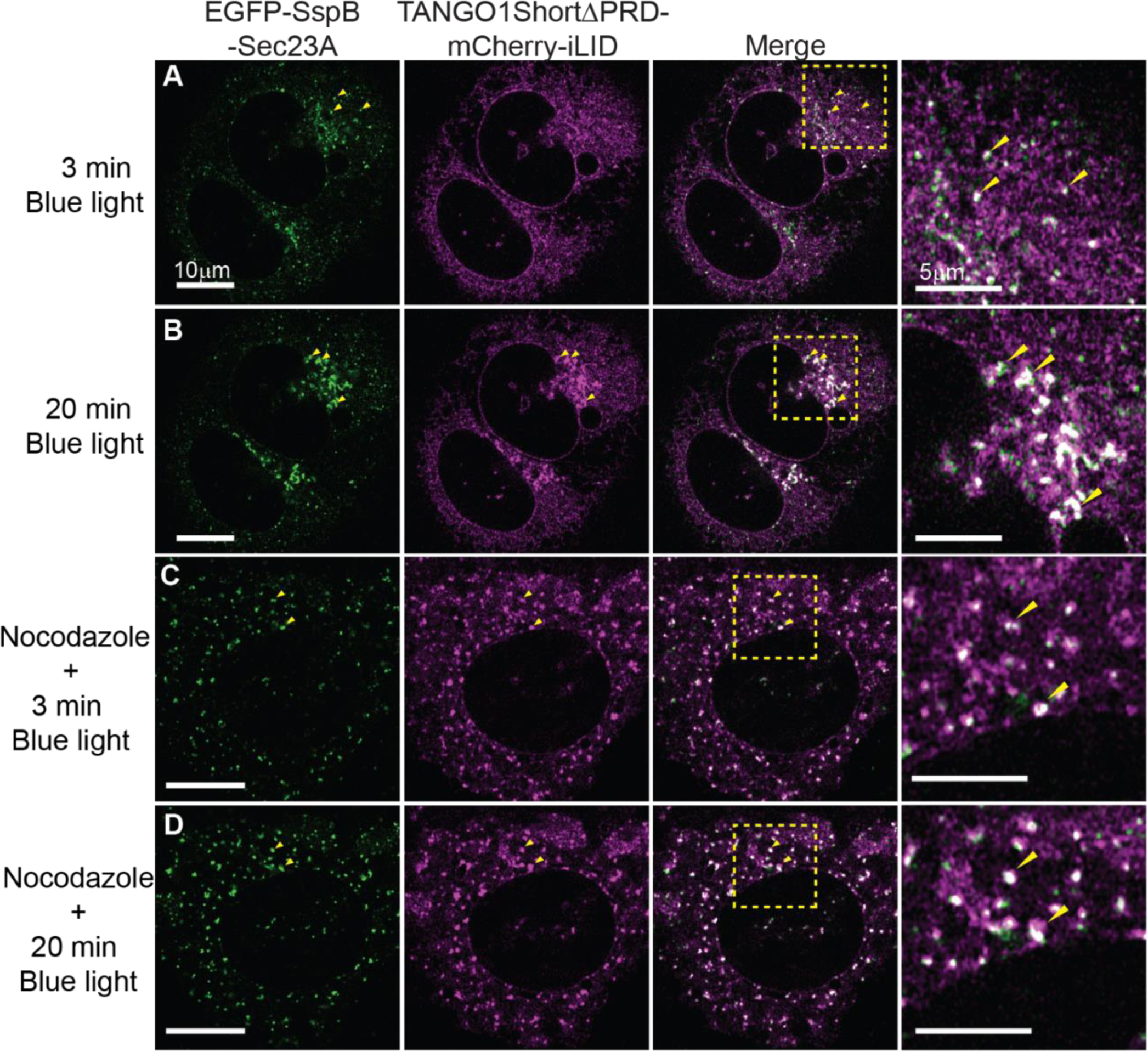
ERES drifts to MTOC upon prolonged TANGO1SΔPRD and Sec23A binding. **(A, B)** Snapshots of timelapse (Supplementary material video 4A*) of U2OS cell co-expressing EGFP-SspB-Sec23A (green) and TANGO1SΔPRD-mCh-iLID (magenta) exposed to blue light for **(A)** 3 min and **(B)** 20min. The time lapse was acquired on Airyscan microscope, scale bar 10 μm. **(A)** ERES marked by EGFP-SspB-Sec23A colocalized with TANGO1SΔPRD-mCh-iLID observed all over the cell, seen as white puncta in overlay image and zoomed image (yellow arrowheads). ERES are heterogeneously spread in the cells, with some of them at periphery and some accumulated at the juxtanuclear region of the cells. **(B)** The same cells at time 20 min show an accumulation of the ERES at juxtanuclear region of the cell. Only few peripheral ERES are visible, whilst the intensity and sizes of the juxtanuclear ERES are significantly higher and observed as fused structures marked by yellow arrowheads in overlay and zoomed images (Scale bar 5 μm). **(C, D)** Snapshots of timelapse (Supplementary material video 4B*) of U2OS cell co-expressing EGFP-SspB-Sec23A (green) and TANGO1SΔPRD-mCh-iLID (magenta) treated with Nocodazole (10.0 μM) for 30 minutes prior to exposure to blue light for **(C)** 3 min and **(D)** 20min. **(C)** ERES marked by EGFP-SspB-Sec23A colocalized with TANGO1SΔPRD-mCh-iLID are observed all over the cell, seen as white puncta in overlay image and zoomed image (yellow arrowheads). ERESs are homogeneously spread in the cells. **(D)** The same cell at 20 min shows no significant change in location of the ERES.

### Prolonged TANGO1SΔPRD-Sec23A interaction interferes with cargo export from ERES

It has been proposed that the sorting at ERES is processive, where the turnover of COPII is utilized to concentrate and sort cargo at ERES.^41^ Previous studies however utilized prolonged expression of mutant Sar1 proteins or in vitro reactions where the GTPase activity of Sar1 was inhibited.^42^ As an alternative approach, we utilized our optogenetic tools to investigate the effects of prolonged (but reversible) TANGO1S-Sec23 binding on cargo secretion. We chose a model cargo, FM4-EGFP-PAUF, a fusion protein of Pancreatic Adenocarcinoma Up-regulated Factor (PAUF) that contains a self-dimerizing domain, DmrD causing its aggregation^43^. When expressed in cells, it is accumulated in the ER (Supplementary Figure 4A). Upon adding a D/D solubilizer, the aggregates dissociate, and the soluble cargo moves to the Golgi within 20 minutes (Supplementary Figure 4B). We combined this synchronized transport of FM4-EGFP-PAUF with the optogenetic tool. FM4-EGFP-PAUF in its aggregated state labels the ER network (**Figure 5A** marked by yellow arrowheads). As previously shown, continuous irradiation with blue light for 20 min, leads to clustering of ERES to the juxtanuclear region (**Figure 4B**), however, this does not affect cargoes trapped in the ER (**Figure 5B**). The mean Pearson correlation coefficient (PCC) of TANGO1SΔPRD-mCh-iLID and FM4-EGFP-PAUF was measured at juxtanuclear region marked by yellow dotted box (details in methods and materials). The PCC shows no statistical difference with time in absence of D/D solubilizer, as the FM4-EGFP-PAUF remains in the ER (**Figure 5C**, empty bars; also see Supplementary Figure 4C). Upon removal of blue light, TANGO1SΔPRD-mCh-iLID and SspB-Sec23A dissociate, and within few minutes TANGO1SΔPRD-mCh-iLID dispersed in the ER membrane (**Figure 5D**). Continuing the same experiment (in the same cell) D/D solubilizer (1.0 μM) was added and immediately cell was exposed again to blue light for 20 min. The D/D solubilizer dissociates the aggregates of FM4-EGFP-PAUF, and the soluble form is available for secretion from the ER (Supplementary Figure 4B). However, upon stabilization of TANGO1S-Sec23 by blue light, the now solubilized FM4-EGFP-PAUF accumulated at ERES that clustered at the juxtanuclear region (**Figure 5E**). Upon TANGO1S-Sec23 dissociation (turning off the blue light), the soluble FM4-EGFP-PAUF left the ER and was trafficked to the Golgi (**Figure 5F**). A significant change in mean of PCC between TANGO1SΔPRD-mCh-iLID and FM4-EGFP-PAUF was evident for the time points 15 min and 20 min (**Figure 5C**, filled bars, also see Supplementary Figure 4D), suggesting that the selective concentration of traffic-competent, disaggregated FM4-EGFP-PAUF at ERES, progressed over time. This contrasts with the lack of concentration of traffic-incompetent aggregated FM4-EGFP-PAUF at ERES during such experiments (**Figure 5B**). Prolonged stable binding of TANGO1S to Sec23 results in a block in secretion of cargo from ER, concentrating the cargo at ERES. Thus, processive activity of the coat inner layer was not required for concentrating FM4-EGFP-PAUF at ERES but blocked ER exit.

**Figure 5:**
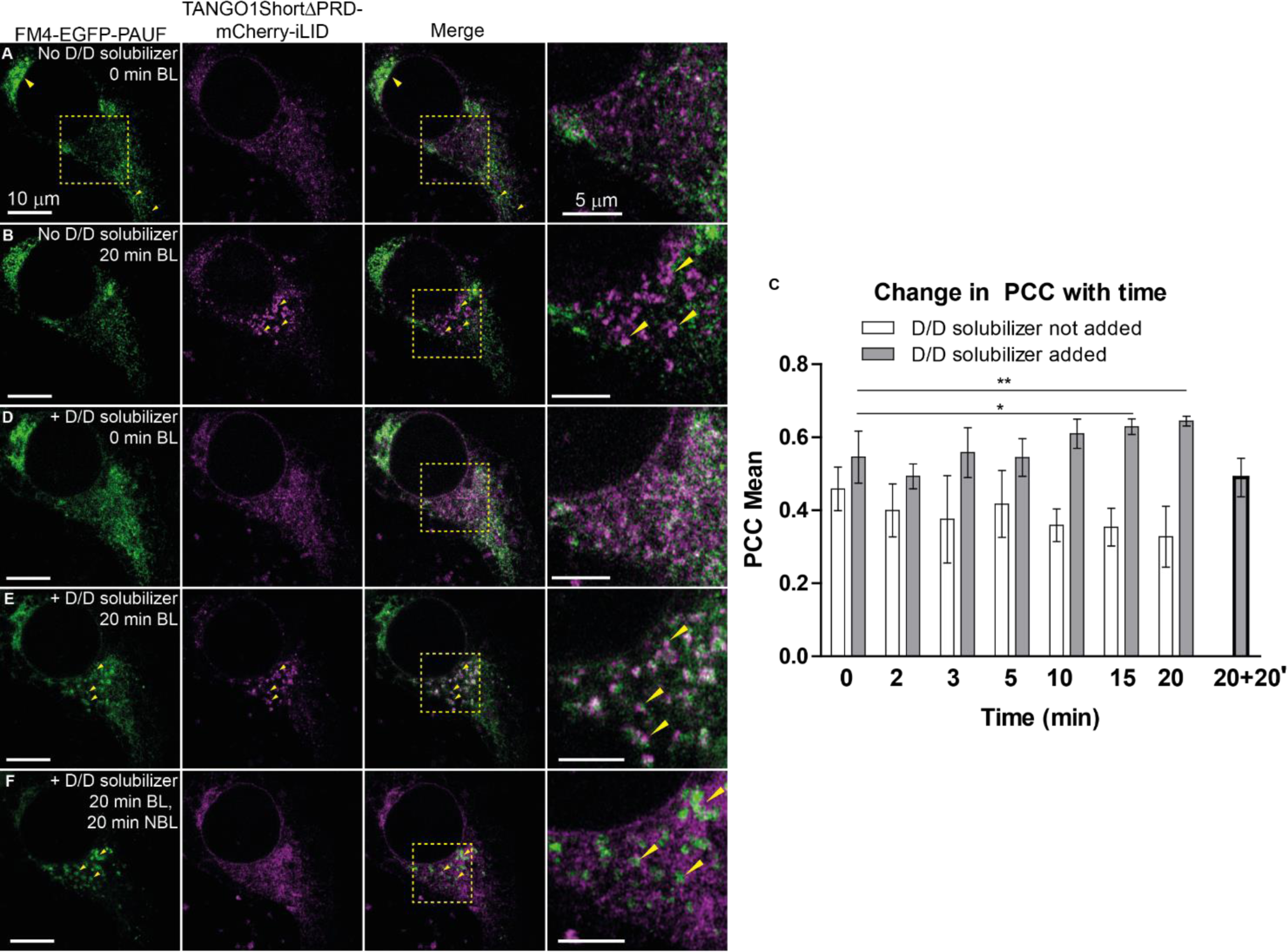
Stable Sec23-TANGO1SΔPRD interaction interferes with cargoes export from ERES. **(A, B)** Snapshots from timelapse (Supplementary material video 5A*) of U2OS cell, stably co-expressing SspB-Sec23A and TANGO1SΔPRD-mCh-iLID, transfected with FM4-EGFP-PAUF. Cells were exposed to blue light for (A) 0 min and (B) 20 min. **(A)** FM4-EGFP-PAUF is localized in the endoplasmic reticulum and in small puncta (marked by yellow arrow heads). The region marked in yellow box shows Golgi region which is devoid of FM4-EGFP-PAUF. The TANGO1SΔPRD-mCh-iLID is present in the ER membrane. **(B)** The same cell exposed to blue light for 20 minutes shows no effect on localization of FM4-EGFP-PAUF. TANGO1SΔPRD-mCh-iLID is now observed in puncta accumulating near the nuclear region of cell (marked by yellow arrowheads), but FM4-EGFP-PAUF is not enriched at ERES. **(C)** A plot of mean of Pearson correlation coefficient (PCC) of TANGO1SΔPRD-mCh-iLID and FM4-EGFP-PAUF against time without (empty bars) and with (filled grey bars) D/D solubilize added. When D/D solubilizer was not added, no significant change in the PCC was observed during the 20 min. One sample t-test against theoretical mean was applied and p value was not significant across the data points. When D/D solubilizer was added, the mean of PCC at time points 15 and 20 min shows significant difference. The time point of cell exposed to blue light for 20 min followed by no light exposure for another 20 min, is marked as 20 + 20’ (bold grey bar). The mean of this time point is not statistically different from the time point zero. The number of asterisks indicate the statistical significance as * for p < 0.05, ** for p < 0.01. (Details in method and materials). **(D, E, F)** (Supplementary material video 5B*) D/D solubilizer (1.0 μM) was added to cells and immediately cells were exposed to blue light for **(D)** 0 min and **(E)** 20 min. **(F)** Image was acquired further after 20 min without blue light exposure. Scale bar 10 μm. **(D)** The same cell in **(B)** was further incubated without blue light for 20 min. FM4-EGFP-PAUF is localized to the ER. TANGO1SΔPRD-mCh-iLID reversibly localizes to the ER membrane. **(E)** D/D solubilizer (1.0 μM) is added and cells were exposed to blue light for 20 min. FM4-EGFP-PAUF is observed in ER as well as accumulating in puncta near the nuclear regions. These puncta are localized with TANGO1SΔPRD-mCh-iLID (marked by yellow arrowheads). **(F)** The reversible dissociation of TANGO1SΔPRD-mCh-iLID and SspB-Sec23A causes dispersion of TANGO1SΔPRD-mCh-iLID into the ER membrane. FM4-EGFP-PAUF’s signal intensity is reduced in the ER and accumulates in vesicular structures in the juxtanuclear region in 20 min.

### Anterograde transport of ERGIC53 is blocked but retrograde transport is unaffected by prolonged TANGO1SΔPRD-Sec23A binding

The lectin ERGIC53 is a transmembrane cargo receptor that cycles between the ER and ERGIC and is predominantly visible as tubulovesicular clusters near Golgi, in the juxtanuclear region (shown in zoom, **Figure 6A**). It moves to ER via retrograde transport and leaves the ER via ERES, hence is observed additionally in the ER (**Figure 6A**). ERGIC53-GFP localizes at ERES labelled by TANGO1SΔPRD-mCh-iLID/SspB-Sec23A upon blue light exposure (**Figure 6B**, marked by yellow arrowheads). Upon continuous irradiation with blue light for 20 min, the ERES moves to the juxtanuclear region. ERGIC53-GFP remains at these ‘clustered ERES’ and accumulates at the juxtanuclear region as well (**Figure 6C**, marked by yellow arrowheads). The colocalization of TANGO1SΔPRD-mCh-iLID and ERGIC53-GFP was significant at time 20 min (**Figure 6D**). The reversal of optogenetic binding led to mobilization of ERGIC53-GFP back into tubulovesicular clusters of ERGIC as it separated from ERES, and simultaneously TANGO1SΔPRD-mCh-iLID dispersed in the ER membrane (Supplementary Figure 5A).

**Figure 6:**
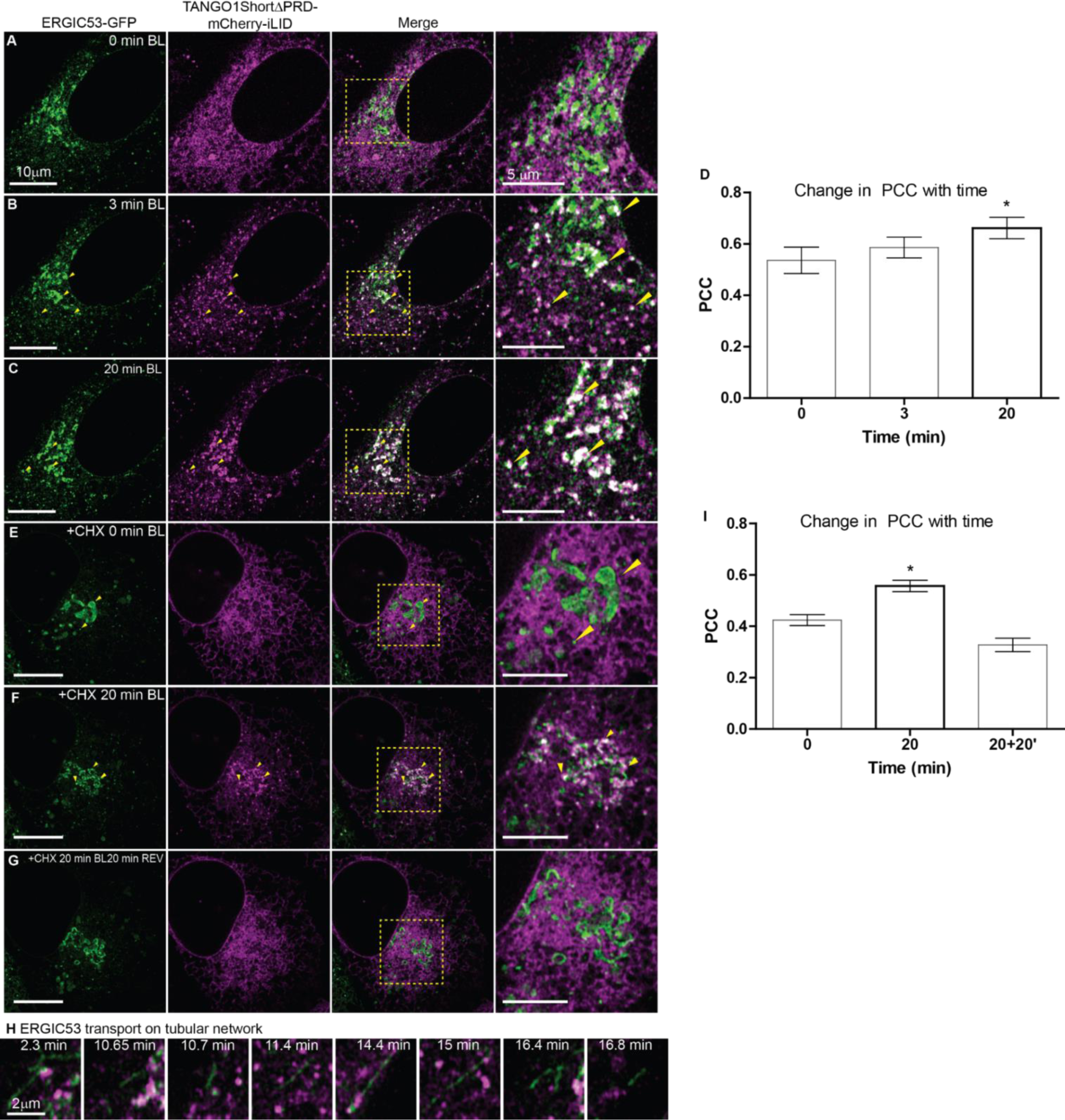
Anterograde transport of ERGIC53 is blocked but retrograde transport unaffected with blue light. **(A, B, C)** Snapshots from timelapse (Supplementary material video 6A*) of U2OS cell, stably co-expressing SspB-Sec23A and TANGO1SΔPRD-mCh-iLID, transfected with ERGIC53-GFP. Cells were exposed to blue light for **(A)** 0 min, **(B)** 3 min and **(C)** 20 min. The time lapse was acquired on Airyscan microscope showing ERGIC53-GFP (green) and TANGO1SΔPRD-mCh-iLID (magenta), scale bar 10 μm. **(A)** ERGIC53-GFP is localized predominantly at juxtanuclear region (Golgi region), and discrete puncta throughout the cell. In the zoom, ERGIC53-GFP localized in the juxtanuclear region is visible as tubulovesicular clusters. TANGO1SΔPRD-mCh-iLID spread in the ER membrane. **(B)** The same cells after 3 min blue light exposure. More discrete puncta of ERGIC53-GFP are observed in the cell. TANGO1SΔPRD-mCh-iLID is seen at discrete puncta (ERES), colocalized with ERGIC53-GFP (seen as white puncta marked by yellow arrowheads). **(C)** The same cell further exposed to blue light for 20 min. ERGIC53-GFP is displaced from its location at Golgi region to puncta (ERES) marked by TANGO1SΔPRD-mCh-iLID. These puncta are accumulated in the juxtanuclear region. The peripheral ERES are also observed colocalized with ERGIC53-GFP and TANGO1SΔPRD-mCh-iLID. (Marked by yellow arrowheads in zoom and merge). **(D)** A plot of mean of Pearson correlation coefficient (PCC) of TANGO1SΔPRD-mCh-iLID and ERGIC53-GFP at time 0, 3 and 20 min of exposure to blue light. One sample t-test against theoretical mean applied, significant change in the PCC was observed during the 20 min. The number of asterisks indicate the statistical significance as * for p < 0.05. (Details in method and materials). **(E, F, G)** Snapshots from timelapse (Supplementary material video 6B*) of U2OS cell stably co-expressing SspB-Sec23A and TANGO1SΔPRD-mCh-iLID (magenta) transfected with ERGIC53-GFP (green). The cells were incubated with cycloheximide (100 μM, for 30 min) prior to exposing to blue light for **(E)** 0 min, **(F)** 20 min and **(G)** 20 min blue light exposure followed by 20 min no blue light exposure. **(H)** Snapshots from timelapse (Supplementary material video 6A*) of U2OS cell stably co-expressing SspB-Sec23A and TANGO1SΔPRD-mCh-iLID (magenta) transfected with ERGIC53-GFP (green) and exposed to blue light up to 20 minutes. Occasionally ERGIC53-GFP can be observed in tubular structures moving away from the juxtanuclear region. **(I)** A plot of mean of Pearson correlation coefficient (PCC) of TANGO1SΔPRD-mCh-iLID and ERGIC53-GFP at time 0, 20 min of exposure to blue light and 20+20’ (20 min blue light + 20 min no blue light) time point. One sample t-test against theoretical mean applied, significant change in the PCC was observed at 20 min (p<0.05). The number of asterisks indicate the statistical significance as * for p < 0.05. (Details in method and materials).

The displacement of ERGIC53-GFP from its juxtanuclear localization to ERES in presence of blue light may reflect retrieval of ERGIC53-GFP from ERGIC followed by its arrest and concentration at ERES. Alternatively, it can be argued that it is the newly synthesized ERGIC53-GFP that concentrates at ERES in our experiments. To test this, cells were treated with cycloheximide to block new synthesis of ERGIC53-GFP prior to exposure to blue light. At time zero, ERGIC53-GFP was observed in its juxtanuclear region and in puncta (**Figure 6E**, marked by yellow arrowheads). Upon exposure to blue light, the juxtanuclear organization of ERGIC53-GFP appeared fenestrated and the ERES were enriched with ERGIC53-GFP (**Figure 6F**, marked by yellow arrowheads). This indicates that ERGIC53-GFP has moved into the ER and its retrograde transport is not affected. However, ERGIC53-GFP was then unable to leave the ER and accumulated at ERES.

When TANGO1SΔPRD-mCh-iLID and SspB-Sec23A dissociate upon removal of blue light, ERGIC53-GFP reassembled into tubulovesicular structure (**Figure 6G**). At the same time, disassembly, and reformation of ERGIC53-GFP tubulovesicular clusters shows that the forward transport at ERES is temporarily blocked and it resumes after blue light is turned off. Occasionally ERGIC53-GFP can be observed in tubular structures moving away from the juxtanuclear region (**Figure 6H**), a phenomenon earlier reported in ER-ERGIC-Golgi recycling of expressed GFP-ERGIC-53 in Hela cells.^44^ The colocalization of TANGO1SΔPRD-mCh-iLID and ERGIC53-GFP in cells treated with cycloheximide, became significant at the 20 min time point as expected, and co-localization was reduced after subsequent 20 min chase with no illumination (**Figure 6I**, mean, standard deviation, p-values, and t-statistics are provided in Supplementary Figure 5B,C). The current observations confirm that retrograde transport of ERGIC53-GFP is unaffected and yet its anterograde transport is blocked during light-activated binding of ERES components. Overall, a pronounced effect was observed on anterograde traffic of ERGIC53-GFP. We further investigated the effects of prolonged binding of TANGO1S and Sec23 on dynamics of Golgi enzymes.

Golgi resident protein, mannosidase II-GFP, was predominantly found localized in cisternae at juxtanuclear region of the cell (**Figure 7A**, marked by yellow arrowheads). Exposure to blue light (20 min) led to clustering of the ERES near this region, closely aligned with the cisternae of mannosidase II-GFP (**Figure 7B**). Prolonging the exposure to 40 min did not affect mannosidase II-GFP localization (**Figure 7C**). Golgi-localization of mannosidase II-GFP remained stable throughout the period of light-activation. The proximity of TANGO1SΔPRD-mCh-iLID and mannosidase II-GFP increased threefold at the 20 min timepoint and remained heightened thereafter (**Figure 7D**). In cycloheximide treated cells (**Figure 7E**), mannosidase II-GFP localization was largely unaffected with 40 min blue light exposure (**Figure 7F**). The apposition of TANGO1SΔPRD-mCh-iLID and mannosidase II-GFP was increased twofold (**Figure 7G**) and lost when illumination ceased (Supplementary Figure 6A). The changes in morphology of mannosidase II-GFP, in cycloheximide treated cells post 40 min of blue light exposure, suggested partial disassembly of Golgi complex (**Figure 7F**). The mild Golgi fragmentation and its apposition to ERES is consistent with Golgi fragmentation associated with inhibition in trafficking, but unlike the fusion and complete relocation of Golgi membranes to the ER as seen with BFA treatment.^45, 46^

**Figure 7:**
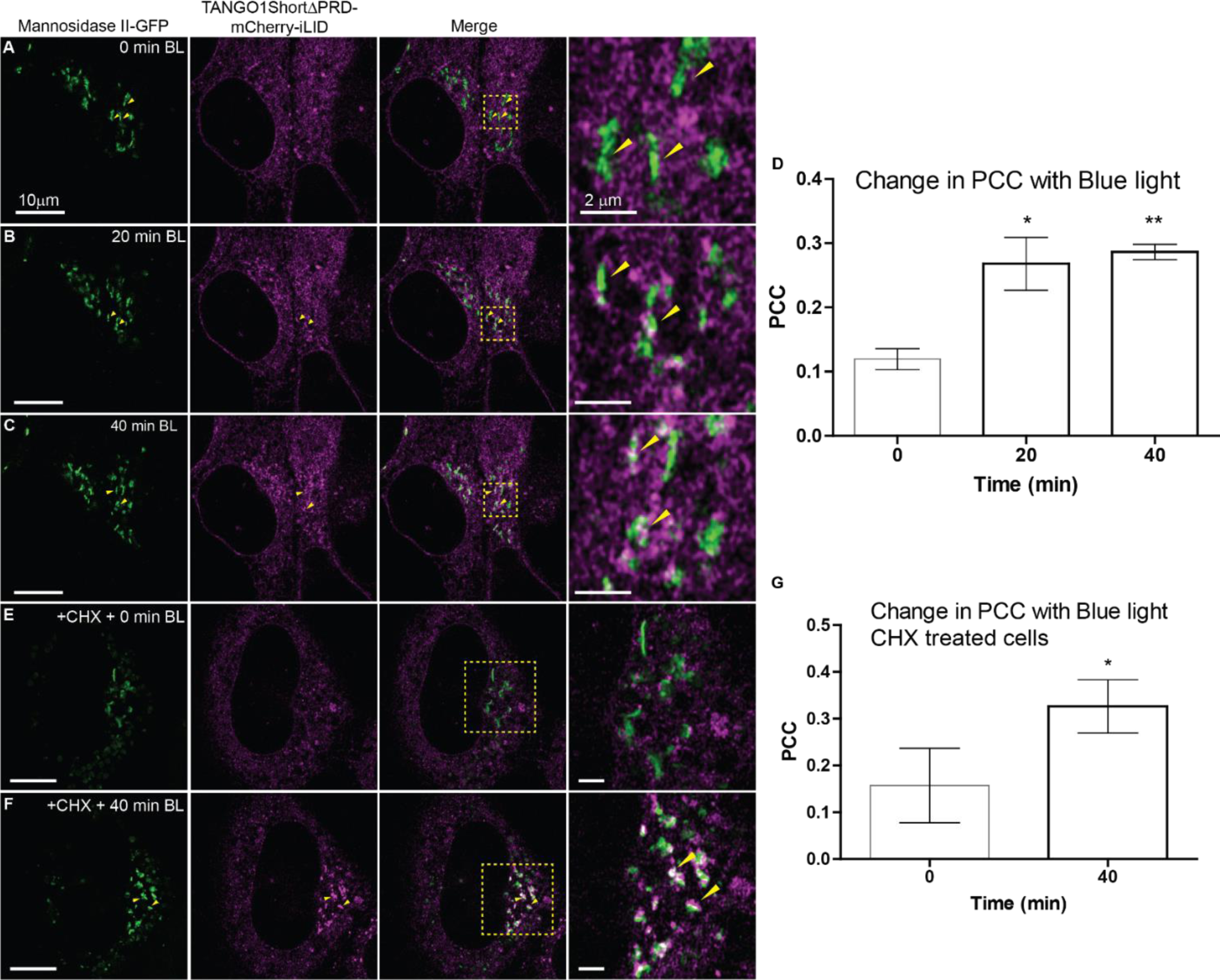
Limited impact on Mannosidase II organization with prolonged Sec23A-TANGO1S binding. **(A, B, C)** Snapshots from timelapse (Supplementary material video 7A*) of U2OS cells, stably co-expressing SspB-Sec23A and TANGO1SΔPRD-mCh-iLID, transfected with mannosidase-II-GFP. Cells were exposed to blue light for **(A)** 0 min, **(B)** 20 min and **(C)** 40 min. The time lapse was acquired on Airyscan microscope showing Mannosidase-II-GFP (green) and TANGO1SΔPRD-mCh-iLID (magenta). Scale bar 10 μm. **(A)** Mannosidase-II-GFP is observed as tubular structures in the juxtanuclear region. (Marked by yellow arrowheads, in zoom and merge). TANGO1SΔPRD-mCh-iLID spread in the ER membrane. **(B)** The same cells exposed to blue light for 20 min. Mannosidase-II-GFP is not displaced and maintain its tubular organization. TANGO1SΔPRD-mCh-iLID is accumulated at ERESs clustered at the juxtanuclear region. The ERES are observed in close contact with cisternae of mannosidase II (Marked by yellow arrowheads, in zoom and merge). **(C)** The same cell further exposed to blue light for another 20 min (total 40 min). Mannosidase-II-GFP organization remains tubular, and cisternae are observed in the juxtanuclear region. Density of TANGO1SΔPRD-mCh-iLID marked ERES is increased at the juxtanuclear region. The clustered ERES are observed in close contact with cisternae of mannosidase II (Marked by yellow arrowheads, in zoom and merge). **(D)** A plot of mean of Pearson correlation coefficient (PCC) of TANGO1SΔPRD-mCh-iLID and mannosidase II-GFP at time 0, 20 and 40 min of exposure to blue light. One sample t-test against theoretical mean applied, significant change in the PCC was observed during the 20 min (p<0.05) and at 40 min (p<0.005). The number of asterisks indicate the statistical significance as * for p < 0.05, ** for p < 0.01. (Details in method and materials). **(E, F)** Snapshots from timelapse (Supplementary material video 7B*) of U2OS cells, stably co-expressing SspB-Sec23A and TANGO1SΔPRD-mCh-iLID, transfected with mannosidase-II-GFP. Cells were treated with cycloheximide (100 μM, for 30 min) prior to exposing to blue light for **(E)** 0 min and **(F)** 40 min. The time lapse was acquired on Airyscan microscope showing Mannosidase-II-GFP (green) and TANGO1SΔPRD-mCh-iLID (magenta). Scale bar 10 μm. **(G)** A plot of mean of Pearson correlation coefficient (PCC) of TANGO1SΔPRD-mCh-iLID and mannosidase II-GFP in cycloheximide treated cell exposed to blue light for 0, and 40 min, and exposed to blue light and no blue light for 40 min each (40+40’). One sample t-test against theoretical mean applied, significant change in the PCC was observed during the 40 min (p<0.05). The number of asterisks indicate the statistical significance as * for p < 0.05. (Details in method and materials).

Overall, the stable interactions between TANGO1S to Sec23 at ERES: (i) prompted microtubule-dependent movement of ERES to MTOC, (ii) hindered ER export of FM4-PAUF-EGFP, (iii) blocked anterograde transport of ERGIC53, and (iv) partially fragmented Golgi complex. Subsequent investigations were carried out to explore the impact of these events on trafficking of cargoes of varying sizes.

### Cargo selectivity at ERES: small size cargo ss-GFP exits from majority of ERES whilst collagen only engage a fraction of ERES

It has been proposed that ERGIC membranes are recruited to ERES by TANGO1 family of proteins for secretion of large cargoes like collagen.^47^ Since prolonged association of TANGO1S and Sec23A causes block of cargo export and alteration in ERGIC53 dynamics (**Figure 5 and 6**), we monitored the secretion of cargoes of different sizes to determine if they behave differently at ERES. We chose ss-GFP as a representative small cargo, likely lacking an exit signal or interactions with dedicated cargo receptors, to study its trafficking at ERES during blue light induced block. At steady state, ss-GFP was observed in the ER and Golgi region (**Figure 8A**). Upon blue light exposure, when TANGO1SΔPRD-mCh-iLID accumulates at the ERES, ss-GFP is found localized with majority of ERES (**Figure 8B, C;** showing representative ERES with and without ss-GFP, see also Supplementary Figure 7A).

**Figure 8:**
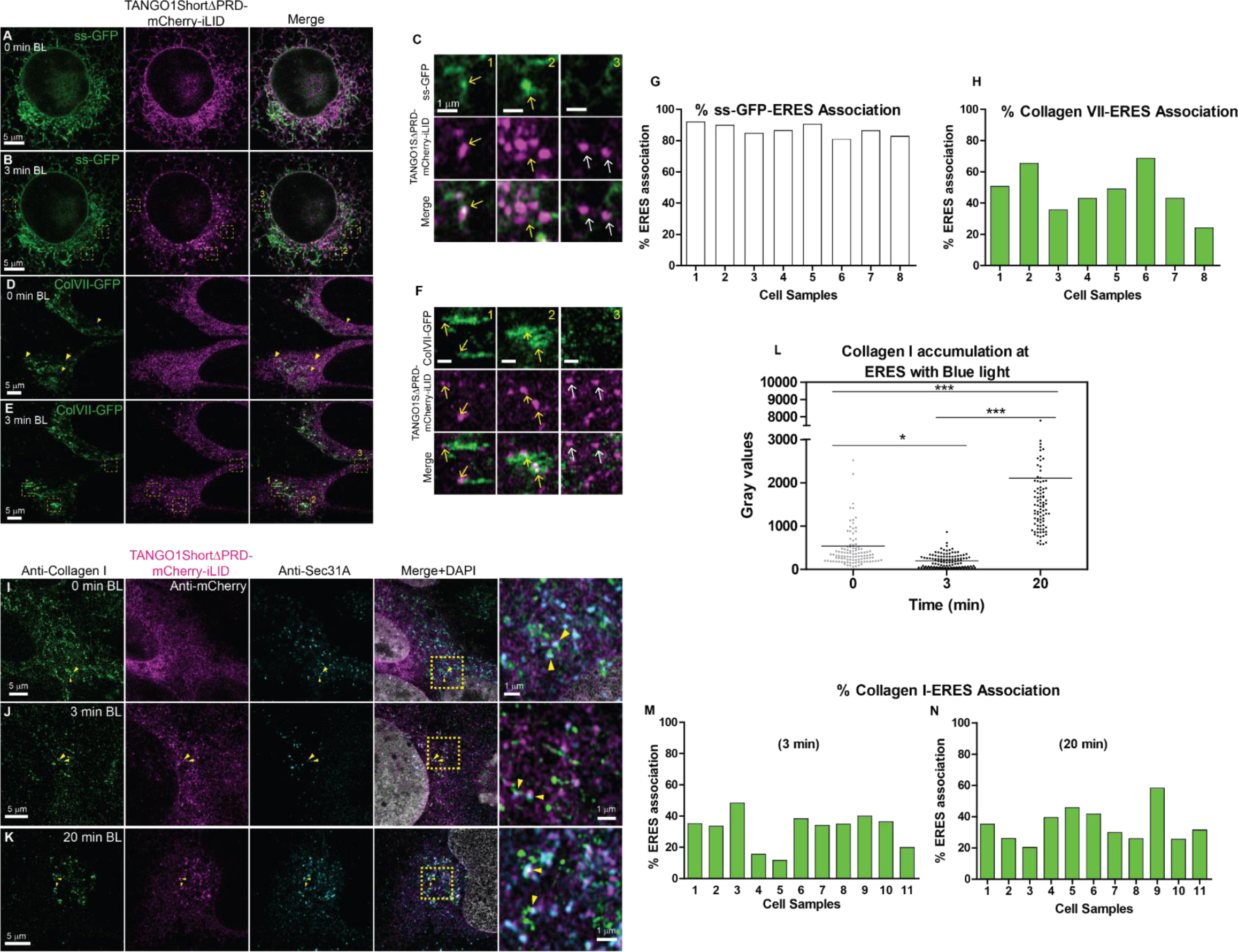
Cargo selectivity at ERES: small size cargo ss-GFP exits from majority of ERES whilst Collagen only engage a fraction of ERES. Snapshots from timelapse (Supplementary material video 8A*) of U2OS cell stably co-expressing SspB-Sec23A and TANGO1SΔPRD-mCh-iLID and transfected with ss-GFP. Cells exposed to blue light for **(A)** 0 min and **(B)** 3 min. Scale bar 5 μm. **(A)** ss-GFP is observed in the ER and Golgi region. TANGO1SΔPRD-mCh-iLID is spread in the ER membrane. **(B)** TANGO1SΔPRD-mCh-iLID is observed in discrete puncta throughout the cell. **(C)** Representative ERESs colocalized with ss-GFP (**1, 2** marked by yellow arrows) and not colocalized with ss-GFP (**3** marked by white arrows) are shown in zoom. Snapshots from timelapse (Supplementary material video 8B*) of U2OS cell stably co-expressing SspB-Sec23A and TANGO1SΔPRD-mCh-iLID and transfected with Collagen VII-GFP. Cells exposed to blue light for **(D)** 0 min and **(E)** 3 min. Scale bar 5 μm. **(D)** Collagen VII-GFP is expressed in the cells as puncta and fibrillar structures (marked by yellow arrowheads). TANGO1SΔPRD-mCh-iLID is spread in the ER membrane. **(E)** Collagen VII-GFP is observed colocalized at the ERES marked by TANGO1SΔPRD-mCh-iLID. **(F)** Shown in zoom are tubular structure of collagen VII-GFP connected to, as well as colocalized with ERES (**1, 2** marked by yellow arrows). Some ERES are observed not localized with collagen VII-GFP (**3** marked by white arrows). **(G, H)** A plot of percentage association of **(G)** ss-GFP and **(H)** collagen VII-GFP with ERES marked with TANGO1SΔPRD-mCh-iLID, upon exposing cells for 3 min with blue light. Total number of cells =11, independent experiments > 3. **(I, J, K)** Microscopy image of U2OS cell stably co-expressing TANGO1SΔPRD-mCh-iLID (magenta) and SspB-Sec23A, irradiated with blue light for **(I)** 0, **(J)** 3 and **(K)** 20min. Cells were fixed and immuno-stained with antibodies against endogenous collagen I and Sec31A. **(I)** Collagen I appear in tubular and puncta form in cell and appears colocalized with ERES marked by Sec31A (Zoom, marked by yellow arrow heads). TANGO1SΔPRD-mCh-iLID is spread in the ER. **(J)** TANGO1SΔPRD-mCh-iLID localizes at the ERES (marked by Sec31A) after irradiation with blue light for 3 min, and collagen I appears closely associated with these ERES (Zoom, marked by yellow arrow heads). **(K)** TANGO1SΔPRD-mCh-iLID and Sec31A marked ERES accumulate in the juxtanuclear region of the cell, and higher concentration of ERES are observed localized with collagen I (Zoom, marked by yellow arrow heads). **(L)** A plot of change in gray values (intensity) of endogenous collagen I associated with ERES marked by Sec31A upon exposure to blue light for 0, 3 and 20 min. A significant decrease in gray value is observed from 0 to 3 min (p<0.05), and a significant increase in gray values is observed at 20 min (p<0.001). The number of asterisks indicate the statistical significance as * for p < 0.05, ** for p < 0.01, *** for p<0.001. One-way Anova test with Tukey’s Multiple Comparison Test in each pair was applied. **(M, N)** A plot of percentage association of endogenous Collagen I with ERES marked with Sec31A, during exposure of cells to blue light at time **(O)** 3 min and **(P)** 20 min. n=11.

Collagen VII is large fibrillar secretory cargo and at steady state, when expressed exogenously in stable cells, it was present in the ER and Golgi and is visible as puncta and tubules (**Figure 8D**). Upon exposure to blue light, collagen VII-GFP was observed in tubular structures adjacent to ERES (**Figure 8E, F,** showing representative ERES with and without collagen VII-GFP). Cargo-ERES association analysis in ∼2000 ERES revealed that ss-GFP exhibited significant association (>80%) whereas, in an independent and similar analysis, collagen VII-GFP showed a restricted association (<50% on average) with ERES (n=8 cells, **Figure 8G, H**, see also Supplementary Figure 7B, C). Analysis of the ERES in contact with collagen VII-GFP further revealed the highly dynamic nature of the tubules that remain bound to the ERES (Supplementary Figure 7D). The size of this tubular collagen VII-GFP were observed ∼ 1μm (Supplementary Figure 7E).

Stable cells (U2OS expressing SspB-Sec23A and TANGO1SΔPRD-mCh-iLID), were exposed to blue light for varying times (0-20 min). Fixed-cell images revealed an association of endogenous collagen I with ERES (marked by Sec31A) at time 0-, 3- and 20-min. Collagen I appeared at few ERES at 0 min (**Figure 8I**) and at 3min (**Figure 8J**, marked by yellow arrowheads).While the ERES enriched in TANGO1SΔPRD-mCh-iLID accumulated at the juxtanuclear region of the cell, clusters of endogenous collagen I also appeared at the same region, co-localized, and in close proximity to ERES (**Figure 8K**). We observed two orders of magnitude increase in the intensity of collagen I at ERES at 20 min time point (**Figure 8L**). Despite the increase in the gray values, collagen I association with ERES exhibited a significant degree of selectivity as it still concentrated at a subset of ERES. The collagen I-ERES association remained constant with time: ∼ 31% at time 3 min (**Figure 8M**), and ∼ 34% at time 20 min (**Figure 8N**). This demonstrated the persistent selective interaction of collagen I with ERES with prolonged blue light exposure.

Occasionally, tubules of endogenous collagen I associated with ERES were also observed, and they displayed two distinct segments. These segments can be classified as segment A, tubule and co-localized ERES and segment B, tubule not co-localized ERES (marked by white and yellow arrowhead respectively, Supplementary Figure 8). The collagen I-ERES associations were investigated for the presence of other proteins in both these segments (Supplementary Figure 8 A-D). The segment A (ERES co-localized collagen I tubule) was co-localized with Sec31 (Supplementary Figure 8A) and ERGIC 53 (Supplementary Figure 8B). However, segment B occasionally lacked ER resident protein, calreticulin (Supplementary Figure 8C), and the chaperone of collagen, HSP47 (Supplementary Figure 8D), indicating that segment B of these tubules might not be a part of the ER. Clearly, there was a link between these tubular structures and ERES and given the exit from ER is hindered (which means that collagen I has not left the ER), it is suggested that collagen I was accumulated at a distinct region that separated ERES from the ER. The exact nature and classification of this ERES-ER connecting region needs to be further investigated and is beyond the scope of this study. However, the current block of ER exit and increased accumulation of bulky cargo at ERES was essential to elicit such interactions at ERES. These ERES were marked by components of a physiological ERES, and we demonstrated that there is a difference in accumulation of small vs large cargoes as diffused vs tubular accumulation, respectively. Additionally, the persistent accumulation of large cargoes at only a specific subset of ERES shows strong evidence of ERES’ inherent heterogeneity towards cargo collection.

## Discussion

### Optical trapping of ERES

The underlying mechanism of how cargoes are collected at ERES prior to their exit from the ER is not fully understood. Analysis of arrested traffic intermediates at ERES is key in defining steps in traffic. These steps include cargo sorting, packing, and generation of export vehicle appropriate for the size and quantities of secretory cargoes. In vivo, temperature-based traffic blocks, or the expression of sensitive mutants, have been very useful in advancing our understanding, but these methodologies rely on non-physiological conditions that affect multiple cellular pathways, which complicates interpretations. ^48, 49^ Cell-free in vitro assays are useful for such analysis yet are based on non-physiological stoichiometries and loss of spatial proximities of compartments.^50, 51^ In contrast, our optogenetic system permits direct control of the ER exit machinery through the binding of TANGO1S to Sec23A at the ERES (**Figure 1B**). This light-based approach allowed us to control coat assembly rapidly, transiently, and reversibly, while maintaining physiological conditions. TANGO1 family members bind Sec23A via their proline rich domain (PRD). In our system, PRD is removed from TANGO1S and modified versions of TANGO1SΔPRD and Sec23A are generated, each bearing one part of an optogenetic system to control their binding (**Figure 1A**). Exogenously expressed TANGO1SΔPRD-mCh-iLID was dispersed in the ER (**Figure 1C**), since removal of PRD prevented its recruitment to ERES.^21, 47^ Exposure to blue light caused rapid relocation of TANGO1SΔPRD-mCh-iLID to pre-existing ERES marked by EGFP-SspB-Sec23A and enriched with Sec16L, Sec13 and cTAGE5 (**Figure 1 and 2**, Supplementary Figure 2, 3). Removing light activation caused rapid dispersal of TANGO1SΔPRD-mCh-iLID, demonstrating the reversibility of our method (**Figure 1E**). H89 treatment inhibits the recruitment and activation of Sar1 on ER membranes.^37, 38^ In our system, H89 treatment prevented TANGO1SΔPRD-mCh-iLID and EGFP-SspB-Sec23A assembly at ERES upon exposure to blue light (**Figure 3F**). Therefore, optogenetically controlled binding of TANGO1S and Sec23A requires Sar1 activation. Thus, the system reproduces the physiological events of COPII assembly yet inhibits the dynamics of the COPII inner layer at ERES providing an opportunity to understand the impact on cargo export.

### ERES are mobilized to the MTOC during prolonged binding of TANGO1SΔPRD to Sec23A

Microtubule (MT) molecular motors dynein is reported to be associated with ERES and to be involved in mobilization of ER-to-Golgi transport intermediates.^40, 52^ Dynein binds to Sec23 using the p150glued-containing dynactin complex, pulling ERES towards the MTOC. In our optogenetic system, the stabilization of Sec23-TANGO1S binding likely prolongs the dynein recruitment and promotes minus (-) end directed movement of ERES to the juxtanuclear region in a microtubule dependent manner. Since TANGO1 proteins do not leave the ER^21, 26, 27^, our data suggests that COPII-MT interactions mobilize ERES in the plane of the ER membrane. It is possible that when coat dynamics are reduced (which likely occurs when exporting bulky cargoes or when traffic is altered by metabolic demands) mobilization and clustering of ERES (**Figure 4B**), near ERGIC-Golgi membranes, is an adaptive response that facilitates traffic. This would occur via tunnels and tubules given the shorter gap between ERES and the ERGIC-Golgi membranes.^53, 54^

### Stable Sec23-TANGO1SΔPRD interaction blocks cargo export

Sorting and concentration of cargo at ERES precedes ER exit. Sorting is achieved by coat interactions between cargo and cargo receptors. ^55, 56^ Some sorting receptors interact with Sec24 and are subsequently exported along with the outgoing cargoes.^57^ Others, such as TANGO1, remain at ERES.^21^ We have found that prolonged binding between TANGO1SΔPRD and Sec23, stalls cargo export from the ER (**Figure 5**). However, tested cargoes are still concentrated at ERES, regardless of their size. For example, ER bound cargo FM4-EGFP-PAUF is normally released upon dissolution by a cell penetrating D/D solubilizer^43^, and exported to Golgi (Supplementary Figure 4A). However, it remained localized with TANGO1S containing ERES with prolonged exposure to blue light (**Figure 5E**). The release of TANGO1S from Sec23A resumed ER export and FM4-EGFP-PAUF transports to the Golgi (**Figure 5F**).

Stable TANGO1S and Sec23A binding at ERES stalls the exit of ERGIC53-GFP without affecting its retrograde transport, eventually leading to accumulation of ERGIC53-GFP at the ERES (**Figure 6**). However, this block of cargo and cargo receptor at the ERES had only minimal effects on Golgi morphology and location of mannosidase II-GFP, even if the cells were exposed to blue light for longer durations (**Figure 7**). This further supports previous data that the process of retrograde traffic of Golgi membrane components is relatively slow compared to the rapidly recycling ERGIC membranes under physiological conditions. ^58^ Why does binding of Sec23A and TANGO1S in blue light inhibit cargo export? The stable capture of Sec23/Sec24 by TANGO1, likely immobilizes associated cargo and cargo receptors, physically prohibiting their exit while deregulating the GTPase activity of Sar1, further leading to defects in membrane remodeling and fission.

What is the physiological significance of the binding of TANGO1 and its family members to Sec23A? We suggest that the binding is a kinetic brake that cells use to generate an export vehicle appropriate for the size and quantities of secretory cargoes. This kinetic brake becomes especially important, for example, in the case of trafficking of collagens that requires recruitment and fusion of ERGIC membranes.

Previous models for cargo concentration at ERES invoked a processive role for COPII.^41, 42^ In these models, dynamic cycles of binding and release between the coat inner layer and cargo/cargo receptors lead to selective enrichment of traffic-competent cargoes at ERES, whereas proteins that display weaker coat binding are discarded. The model is mainly based on outcomes observed during prolonged expression of COPII mutants and in particular of GTPase deficient Sar1 in cells. Under these conditions some cargoes are concentrated at ERES while others are not.^41, 42, 59^ Our application of optogenetics to control the dynamics of the coat inner layer, shows that cargoes concentrate even in the relatively short period of coat inner layer immobilization. Thus, coat processivity may not be obligatory for concentrating cargoes at ERES.

### Cargo and carrier selectivity at ERES

Our findings have demonstrated that a significant distinct population of ERES handles cargoes of different sizes. These differential associations of cargoes with ERES were accentuated by stabilizing TANGO1S-Sec23A binding (**Figure 8**). While small cargoes associated with the majority of ERES, bulky cargoes like collagen I and VII revealed a more selective accumulation at ERES (**Figure Schematic 9A**). TANGO1 family members are found at ERES, how is the specificity for collagens at less than half the total number of exit sites achieved? It has been reported that levels of Sec16A, Sec24B and Sec31A can vary amongst ERES and their abundance are likely regulated.^60^ Studies on mechanism of polarized basement membrane secretion in Drosophila cells, revealed an increased levels of Tango1 at basal ER exit sites and suggested these ERES are specialized for collagen export.^61^ We propose that it is possible that additional components interact with the TANGO1 machinery for use of some ERES for the export of bulky cargoes. The small cargoes could be exported along with the collagens (by tubules/tunnels) or from other ERES by the standard COPII vesicles. This would explain why depletion of TANGO1 affects predominantly the collagens, but there is also an effect on the exit of other cargoes.^62, 63^ When cells are completely depleted of TANGO1^25^, or TANGO1 is trapped with Sec23A, as in our experiments, all exit sites are affected (**Figure Schematic 9B**). Under these conditions, there is a global defect in cargo export in addition to rapid relocation of ERGIC membranes to the ER.

**Figure Schematic 9:**
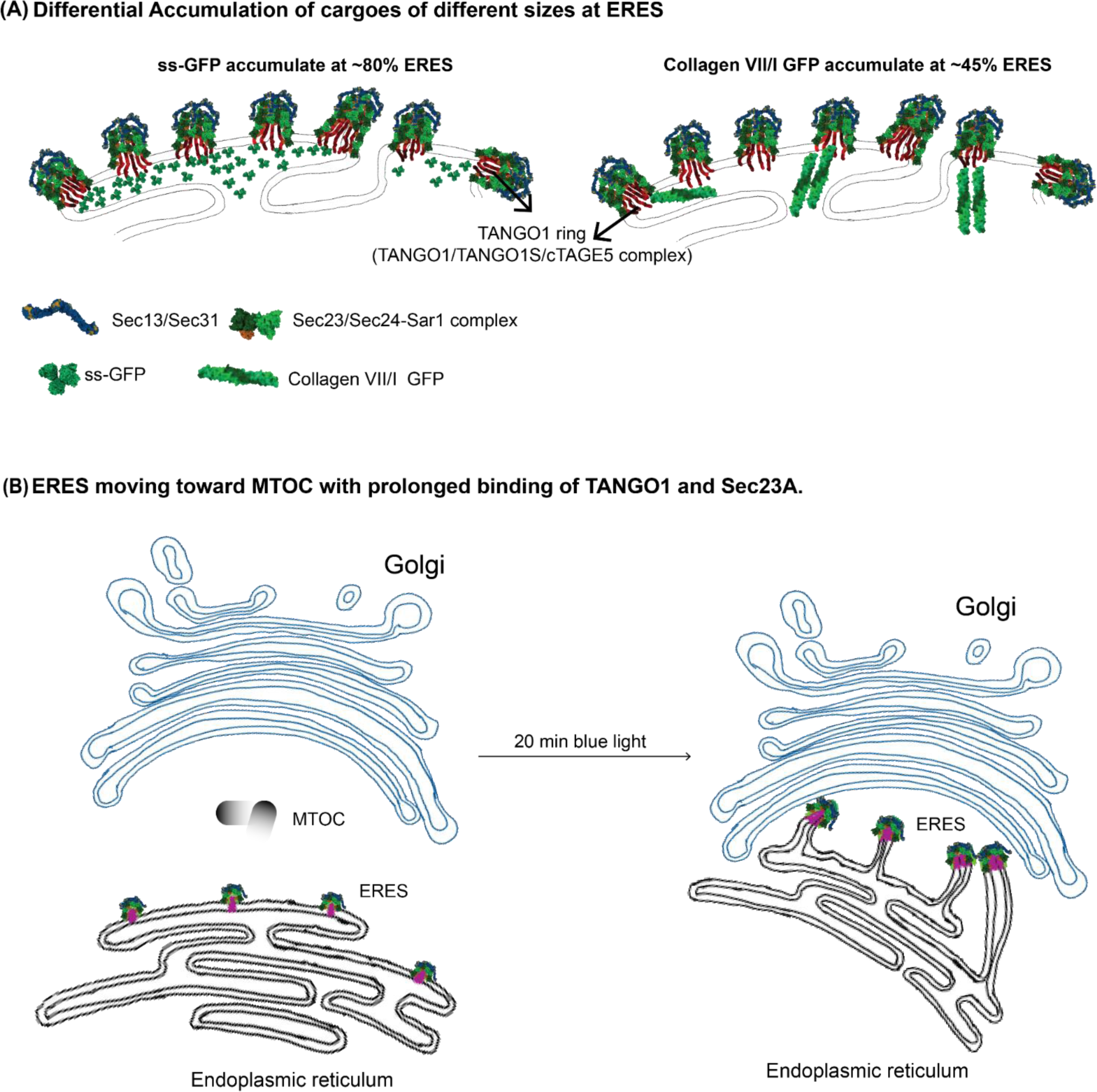
**(A) Schematic of Differential accumulation of ss-GFP and Collagen VII/I-GFP.** While ss-GFP is present at majority of the ERES. Collagen VII/I GFP is restricted to 45% of ERES. **(B) ERES migration to MTOC.** The stable binding of TANGO1 and Sec23A causes migration of ERES in the plane of ER membrane towards the MTOC.

Another important feature of our studies is the appearance of collagen VII-GFP and collagen I in tubular elements close to the ERES. Occasionally, the structures did not colocalize with calreticulin and HSP47 but were observed in proximity of Sec23A and TANGO1S (Supplementary Figure 8). We suggest these features are suggested to be reminiscent of the accumulation of collagen in saccular uncoated domains of the ERES prior to exit where these are segregated from small cargo. The optogenetic arrest of exit likely expanded those domains which are now visible at light resolution. The results support recent studies showing that HSP47 is excluded from ERGIC tubules that mobilize collagens to the Golgi complex and suggest that a pH gradient that facilitates HSP47 release from procollagens is established within ERES. This gradient may facilitate the concentration of collagen at ERES as recently reported. ^64^

We suggest that this mechanism of segregating ERES provides means to a cell to build unique tunnelling mechanism for collagen export while keeping other exit sites open for trafficking of smaller cargoes and to ensure the integrity of the ERGIC compartment. In principle, this would permit cells to use tunnels/tubules for collagen export and small vesicles for smaller cargoes simultaneously. The propensity for the use of these two exit sites would depend on the cell type and physiological demands.

## Materials and Methods

### Antibodies

<u>Primary Antibodies</u> Anti-Collagen I (Abcam, Cat. No.: ab138492), Anti-Sec16A (Sigma, Cat. No.: HPA005684), Anti-Sec31A (BD, Cat. No.: 612350), TANGO1 (MIA3) (Proteintech, Cat. No.: 17481-1-AP), Hsp47 (Enzo, Cat. No.: ADI-SPA-470-F), Calreticulin (Enzo Cat. No.: ADI-SPA-601), ERGIC 53 (Enzo, Cat. No.: ENZ-ABS300-0100), GALNt2 polyclonal (Novus Biological, Cat. No.: AF7507), cTAGE5 (Atlas, Cat. No.: HPA000387), Anti-GFP (Roche, Cat. No.: 11814460001), Anti-RFP[5F8] (Chromotek, Cat. No.: 5f8-20), Anti-mCherry (Origene, Cat. No.: AB0040-200), Anti-calnexin (Abcam, Cat. No.: ab22595). <u>Secondary antibodies:</u> For western blot: a-Rabbit HRP, a-Goat-HRP, a-Rabbit 680.

### Blue-light Activation set up in lab

For fixed cell imaging a Blue light LED-Setup was built in lab. 8 LEDs with wavelength 490 nm emission, were placed on a bread board and attached to an Arduino set up. The Arduino set up was programmed for irradiating blue light 200ms after every 300ms.

### Calculation of Feret Diameter using Image J

Manually visible particles were marked in a region of interest. Feret diameter for each region of interest was obtained using analyze particle in Image J. The mean of Feret particles was plotted against the protein. Feret diameters of 300 distinct puncta were measured in at least three different experiments.

### Cell culture

U2OS cells and the stable cell lines generated were maintained in Dulbecco’s Modified Eagle’s Medium (DMEM, Lonza) cell culture media supplemented with 10% (vol/vol) fetal bovine serum (FBS, heat inactivated, Gibco.). 1% vol/vol penicillin and streptomycin were added to the media. Cells were grown at 37°C, in humidified incubator with 5% CO_2_. As a general practice, all the cells were routinely checked for mycoplasma contamination. Only mycoplasma free cells were used for the experiments.

### Cell fixing and antibody staining

Cells grown on coverslips were washed with 1X PBS 2 times (5 min), fixed with 4% paraformaldehyde (Sigma Cat. No.: P6148) in PBS, followed by washing 3 times for 5 min. Cells were permeabilized using 0.1% Triton-X-100 (Sigma Cat. No.: Cat. No.: T8787) for 10 min and washed 2 times for 5 min. Cells were incubated in blocking buffer 2% BSA for 30 min. Cover slips were mounted on a dry and clean box, and cells were incubated with 50μL primary antibody solution (dilution 1:1000) in 2% BSA (Bovine Serum Albumin, Sigma Cat. No.: A7906) at 4°C cold room overnight. Cells were washed with 1X PBS 3 times (5 min) and incubated in secondary antibody (Dilution 1:1000 in 1X PBS) at RT for 1h in dark. For nuclear staining 0.5µg/µL of DAPI (4’, 6-diamidino-2-phenylindole, dihydro-chloride, Invitrogen. Cat. No.: D3571) was added along with secondary antibody. The cells were washed 3 times with 1X PBS for 5min, in dark. And coverslips were placed on microscopy glass slides using mounting media (FluorSave® reagent, CalBiochem. Cat. No.: 345789). The slides were let to dry at RT for 2h, and stored at 4°C. Prior to imaging the glass slides were brought to RT.

### Cell transfection

For transient transfection X-tremeGENE^TM^ 9 DNA and X-tremeGENE^TM^ HP DNA transfection reagents were used. The ratio of plasmids to reagent was maintained at 1:3, and prepared in Opti:MEM® (Gibco, Cat: # 31985062) media, and incubated at rt for 15 min. The transfection complex was added to the cells (in complete media) slowly, and the plates were swirled gently. Cells were incubated for 24h, and fresh media was supplied to cells to further incubate the cells for 24h.

### ERES-Protein association plots

We quantitatively calculated the differential association of collagen VII-GFP and ss-GFP with the ERES marked by TANGO1SΔPRD-mCh-iLID as representative of cargoes of varying sizes. Associated and non-associated ERES protein events were counted manually for highest accuracy in 8 independent experiments. Using these counts we computed the association of cargoes in each of these 8 cells. To avoid any artifact due to variation in number of ERES, these events were plotted in terms of % occupation of total ERES. For Collagen VII-GFP, efforts were made to choose cells with low expression levels, however, the choice of cells was limited to cells expressing Collagen VII-GFP while still retaining TANGO1ΔPRD-mCh-iLID signal.

### iDimerize Reverse Dimerization System

1.0 μM of D/D solubilizer (Takara, Cat: # 635054) in Opti:MEM® (Gibco, Cat: # 31985062) media was added to cells.

### Microscopy Imaging

Majority of live cell imaging cell were maintained at 37°C in presence of 5% CO_2_ in complete DMEM. Alternatively, complete DMEM was supplemented with 25mM HEPES or Opti:MEM® (Gibco, Cat: # 31985062) media was used directly.

Microscope used:

1. Andor Dragonfly Spinning Disk confocal system (Borealis illumination, 488 and 561 nm lasers)
2. Inverted Leica TCS SP8 confocal microscope (405, 488, 561, 634 nm lasers).
3. Leica STELLARIS confocal system with white light laser (WLL: 485 - 685, 405, 488, 561 nm lasers)
4. Zeiss LSM 980 with Airyscan 2 (405, 488, 561, 639 nm lasers)

High resolution live cell images were acquired on Zeiss LSM 980 with Airyscan 2, in airyscan mode using 32 detectors. Leica Stellaris confocal system was used for H89 treated live cells. Live cell imaging for puncta fusion and fission was recorded on Andor Dragonfly Spinning Disk confocal system (Borealis illumination, 488 and 561 nm lasers).

1. Time lapse acquired on an Andor Dragonfly Spinning Disk confocal system with Borealis illumination, 488_Confocal was used for GFP and 561_Confocal was used for mCherry excitation. Time series was acquired by exciting GFP by 488 nm laser for 200 ms followed by excitation of mCherry by 561 nm laser for 100 ms, and an interval of 200ms was given between each cycle. Total duration of an interval was 500 ms. Drift stabilization was enabled, and iXon EMCCD camera was used capturing images of 512 × 512 pixels (63.40 × 63.32 µm, xy). Time lapse was saved as a sequence of 16-bit pixel depth images. Objective used was 60 X.
2. Fixed cell images on Inverted Leica TCS SP8 confocal microscope and Leica STELLAIRS confocal system. Images were captured using 3 lasers (405, 488, 561 nm lasers). Objective used was 63 X.
3. Fixed cell images on Zeiss LSM 980 with Airyscan 2. 63 X objective was used. Images were acquired on airyscan detector. Bits/pixel: 16.
4. Live cells time lapses were acquired on Zeiss LSM 980 with Airyscan 2 with an airyscan module. Cells were excited using 405, 488, 561 and 633 nm lasers. Objective used: Plan-Apochromat 63X / 1.4 Oil. Videos were acquired and iLID was activated using 488 nm laser (laser power 2), in line scanning mode with no averaging. Airyscan SR was used for optogenetics experiments. Images were acquired at 37°C in a CO_2_ inbuilt chamber. The images and time lapse videos were processed by 2D and 3D airyscan processing in the Zen Software of the microscope.

### Pearson correlation coefficient calculations

To understand the relation between the cargo accumulation at ERES, we employed Pearson correlation coefficient calculations. Given the distinct distribution of TANGO1SΔPRD-mCh-iLID and FM4-EGFP-PAUF, with the former localized predominantly at the ERES (after blue light exposure), and at MTOC region (at 20 min), and the latter predominantly at ER, we tailored our calculations of PCC to only the MTOC region shown in the zoom of the figures. This approach was adopted for specified and relevant assessment of correlation. Hence the analysis was aimed to capture the spatial relation between TANGO1SΔPRD-mCh-iLID and FM4-EGFP-PAUF only in the MTOC region, and the other regions of the cells were excluded from the PCC calculations to avoid dilution in the potential PCC. This approach was extended in calculating the PCC of TANGO1SΔPRD-mCh-iLID with other proteins like ERGIC53-GFP and ManII-GFP as well.

For the timelapse shown in Figure 5, the primary aim was to monitor the effect of blue light on FM4-PAUF-EGFP with and without D/D solubilizer, in the same cell during the different time points. However, due to rapid response to D/D solubilizer to disaggregation of FM4-PAUF-EGFP, monitoring same cell became challenging. Hence the data reported is comprehensive result of:

#### Single-Cell Measurements

The cell where we monitored the response of cargo to D/D solubilizer in the same cell was used for representation in the Figure 5. This allowed for continuous capture of response with time from a single cell.

#### Multiple-Cell Measurements

Since, we were only interested in probing the cargo movement at the ERES we adapted the experiment and acquired the measurement at different time points and extended it to different cells. While this approach might introduce variability due to lack of single cell data acquisition, it was essential to capture the rapid dissolution of FM4-PAUF-EGFP with D/D solubilizer, and its accumulation at the ERES. It was found that the variations in the data sets obtained from multiple cells were not significant. These results were used for measuring the PCC values and have been show in the plots.

For measurement the mean value of PCC at time points 0 was chosen as a reference (hypothetical) point, and a one-sample t-test was applied. Here null hypothesis (*H*_0_) was a mean value equal to the reference value. And alternative hypothesis (*H*_1_) corresponding to the mean value different from the reference value. p-value threshold of 0.05 was used to measure the statistical significance.

### Plasmid generations

1. TANGO1Short-mCherry-iLID was generated from TANGO1Short-FLAG and PEX-mCherry-iLID using in-Fusion cloning. The iLID fragment was originally obtained as a generous gift from the Kuhlman lab (Addgene 60411). TANGO1Short was amplified using 5’-GATCCGCTAGCGCTACCGGTGCCACCATGGACTCAGTACCTGCC-3’ and 5’-ACTACTACCACTACTACCTGGGCTCTGTTTTAAAGCCTG-3’, mCherry-iLID was amplified using 5’-GGTAGTAGTGGTAGTAGTATGGTGAGCAAGGGCGA-3’ and 5’-TCGAAGCTTGAGCTCGAGATCTTTAAAAGTAATTTTCGTCGTTCGCT-3’.The fragments were inserted into a pEGFP-C1 vector (Addgene 46956) using the AgeI/BglII restriction sites.
2. TANGO1ShortΔPRD-mCherry-iLID was generated from TANGO1ShortΔPRD-FLAG and PEX-mCherry-iLID using in-Fusion cloning. The iLID fragment was originally obtained as a generous gift from the Kuhlman lab (Addgene 60411). TANGO1ShortΔPRD was amplified using 5’-GATCCGCTAGCGCTACCGGTGCCACCATGGACTCAGTACCTGCC-3’ and 5’-ACTACTACCACTACTACCTTCTTCTTGCAGCATTGCC-3’, mCherry-iLID was amplified using 5’-GGTAGTAGTGGTAGTAGTATGGTGAGCAAGGGCGA-3’ and 5’-TCGAAGCTTGAGCTCGAGATCTTTAAAAGTAATTTTCGTCGTTCGCT-3’. The fragments were inserted into a pEGFP-C1 vector (Addgene 46956) using the AgeI/BglII restriction sites.
3. EGFP-micro-Sec23A was generated from Sec23A-FLAG and micro-Sec61 using in-Fusion cloning. The SspB (micro) fragment was originally obtained as a generous gift from the Kuhlman lab (Addgene 60410). Sec23A was amplified using 5’-GGTAGTGGTAGTGGTAGTATGACAACCTATTTGGAATTCATTC-3’ and 5’-CGGTACCGTCGACTGCAGTCAAGCAGCACTGGACAC-3’, SspB (micro) was amplified using 5’-CGAGCTGTACAAGTCCGGAAACAGCCGCGTGATGGAATTCAGCTCCCCG-3’ and 5’-ACTACCACTACCACTACCACCAATATTCAGCTCGTCAT-3’. The fragments were inserted into a pEGFP-C1 vector (Addgene 46956) using the PstI/BspEI restriction sites.
4. EGFP-SspB-Sec23A-pHRSIN segment was amplified from Micro-Sec23A-GFP plasmid by PCR using the forward primer: 5’-GAGTCGCCCGGGGGGGaTCCAGATCCGCTAGCGCTACC-3’ and reverse primer:5’TTGCATGCCTGCAGGTCGACTCAAGCAGCACTGGACACAGC-3’. The fragment was inserted in empty plasmid pHRSIN digested by BamHI and SalI, using Gibson assembly.
5. TANGO1ShortΔPRD-mCh-ILID-pHRSIN fragment was amplified from TANGO1ShortΔPRD-mCh-ILID by PCR using the forward primer 5’-GAGTCGCCCGGGGGGGaTCCATGGACTCAGTACCTGCCAC-3’ and the reverse primer 5’-TTGCATGCCTGCAGGTCGACTAAAAGTAATTTTCGTCGTTCGCTGCC-3’, and inserted into empty pHRSIN vector digested by BamHI and SalI using Gibson assembly.
6. TANGO1Short-mCh-iLID-pHRSIN fragment was amplified from TANGO1Short-mCh-iLID by PCR using forward primer 5’-TTGCATGCCTGCAGGTCGACCTTTAAAAGTAATTTTCGTCGTTCGCTG CCT -3’ and reverse primer 5’-GAGTCGCCCGGGGGGGaTCCcaccATGGACTCAGTACCTGC-3’and inserted into polylinker pHRSIN vector digested by BamHI and SalI using Gibson assembly.
7. Micro-Sec23NoGFP was generated by using PCR based site directed mutagenesis using forward primer 5’-GCTAGCGCTACCGGTCGCCACCATGGAATTCAGCTCCCCGAAACGCCC-3’ and reverse primer 5’-GGGCGTTTCGGGGAGCTGAATTCCATGGTGGCGACCGGTAGCGCTAGC-3’. The new fragment generated did not have EGFP and prior to bacterial transfection, the resultant plasmid was subjected to DpnI restriction enzyme.
8. SspB-Sec23A_L309: IRES and PuroR fragements were inserted in L309_EGF plasmid digested by EcoRI and BsrGI using Gibson assembly. ECoRI and BsrGI restriction enzyme sites were introduced in SspB-Sec23A-NoGFP fragment, by first PCR amplifying it from Micro-Sec23NoGFP using forward primer 5’-GCAGCAGGATCCCGCCACCATGGAATTCAGCTCCCC-3’ and reverse primer 5’-GCGGCCGCGCAGCATCAAGCAGCACTGGACACAGCAAG-3’ and inserting it in pcDNA3 vector. Further the SspB-Sec23A-NoGFP (With desired enzyme sites) was PCR amplified using forward primer 5’-AAGCTTGGTACCGAGCGGATCCGCCACCATGGAATTCAGCTCCCC-3’ and reverse primer 5’-GGGGGGGGGGGGCGGAATTCGCTAGCTCAAGCAGCACTGGACACAGCA-3’. This fragment was inserted in the LC309_puromycin vector using Gibson assembly to give desired construct.
9. EGFP-FM4-PAUF: EGFP fragment was amplified from pcDNA3-NtermEGFP using forward primer 5’-ATGGTGAGCAAGGGCGAGGAGC-3’ and the reverse primer 5’-CTTGTACAGCTCGTCCATGCCG-3’. The FM4-PAUF fragment was amplified from pC4S1-ss-mKate2-FM4-PAUF WT (graciously donated by Dr. Yuichi Wakana and Dr. Felix Campelo) using forward primer 5’-GCATGGACGAGCTGTACAAGGGCGCAGCAGCGGGATCTAG-3’ and reverse primer 5’-CACTGTGCTGGATATCTGCAGAATTCCTAGCGACCCACGGGTGAGTTTG-3’. hGH signal sequence from the same plasmid using forward primer 5’-TAGGGAGACCCAAGCTGGCTAGCACCATGGCTACAGGCTCCCGG-3’ and reverse primer 5’-CTCGCCCTTGCTCACCATAAGCTTGGCACTGCCCTCTTG-3’. The three fragments hGH signal sequence, EGFP and FM4-PAUF were inserted in and empty plasmid pcDNA3 by using the NheI and EcoRI restriction enzymes and Gibson assembly.

### Preparation of Stable cell lines

**1.** Generations of stable lines of U2OS co-expressing TANGO1ShortΔPRD-mCh-iLID and EGFP-SspB-Sec23A: Lentiviral particles were generated in HEK-293T cells as described in reference 23. Briefly, TANGO1ShortΔPRD-mCh-ILID-pHRSIN and EGFP-SspB-Sec23A-pHRSIN were separately added to 8× 10^5^ HEK−293T cells with a packaging vector pool (comprising of pCMV 8.91 and pMDG) using TransIT®-293 transfection reagent (Mirus Bio LLC). Post transfection cells were incubated, and viral particles rich supernatant media were harvested after 48h. and filtered through 0.45 µm membrane filter. The viral particles TANGO1ShortΔPRD-mCh-iLID of were added directly to U2OS cells, and the cells were sorted by fluorescence-activated cell sorting (FACS). Subsequently, the resultant stable line expressing TANGO1ShortΔPRD-mCh-iLID, were further treated to viral particles of EGFP-SspB-Sec23A to generate the desired cells using the same protocol.
**2.** U2OS cells stably expressing TANGO1Short-mCh-ILID and EGFP-SspB-Sec23A: were generated using the same approach.
**3.** U2OS cells stably expressing TANGO1ShortΔPRD-mCh-ILID and SspB-Sec23A-NoGFP: Virus particles for SspB-Sec23A-L309 were generated by co-transfecting transfer plasmid (1μg/mL) and pRSV–REV, pMDLg/pRRE, VSV-G 54 (total 1μg/mL) in 300 μL Opti:MEM® media and 6 μL TransIT®-293 in 8 × 10^5^ HEK−293T cells. The cells were incubated for 24h and the media was aspirated, collected, and replaced with fresh media. After another 24h, the media was collected, added to previous aliquot, and filtered through a 0.45 µm membrane filter. U2OS cells stably expressing TANGO1ShortΔPRD-mCh-ILID were seeded in the 60 mm dishes. To one plate, was added 1.0 mL of the viral particles in 2 mL complete media and 25 μL polybrene (10 μg/mL, hexadimethrine bromide, Sigma Cat. No.: 107689). After 24h, the media was replaced with fresh media and the cells were sorted by treating the cells and controlling with puromycin (Gibco Cat. No.: A11138-03). The cells were further tested for successful incorporation of the SspB-Sec23A-NoGFP by microscopy imaging (Supplementary Figure 3A-E).

### Quantification of Number of Puncta Using ImageJ (Trackmate Plugin)

The number of puncta were quantified by adopting a feature of Trackmate plugin in ImageJ.^65^ The image pixel height and width were automatically used by the plugin. Image was filtered using Laplacian of Gaussian (LoG) detector. The estimated blob diameter (size of puncta) was selected in a range of 0.8-1.0 microns. To ensure comprehensive marking of all puncta in each image the threshold was adjusted in the range of 1.0-10.0. The number of puncta were measured in the last frame of a 3 min time lapse of n=15 cells recorded on Airyscan microscope at high resolution. A replicate of minimum 3 or more independent experiments were used for quantifications.

### Western blot

#### Sample Preparation

Cell lysates were prepared from PBS washed cells using lysis buffer containing Tris 50nM, NaCl 150nM, 1mM EDTA (Sigma, Cat. No.: E6758) at pH 7.4 and 1% Triton-X-100 (Sigma, Cat. No.: T8787). Protease inhibitors Leupeptin (5 µM, Focus Biomolecules. Cat. No.: 10-1346), Aprotinin 2 µg/µl (Abcam. Cat. No.: ab146286); Pepstatin A 2 µg/µl (Panreac Química. Cat. No.: A2205) were added to the above. Cells were incubated on ice for 15 min with regular shaking after every 5 min. Cells lysates were centrifuged (12000 RPM, 4 °C,15 min), and supernatant was collected and stored at -20°C.

#### Gel electrophoresis

The lysates with loading buffer (boiled for 10 min), was loaded on 6% polyacrylamide gel, for SDS-PAGE electrophoresis, and ran for 180 min at 80V.

#### Transfer

The proteins were transferred to PVDF membrane, 0.45 μm (Amersham. Cat. No.: 10600023) for 180 min at 80 V. The transfer was carried out in an ice-bucket. The membranes were blocked using 2.5% non-fat dry milk for 30 min at RT.

#### Primary antibody incubation

The membranes were treated to primary antibodies at a dilution of 1:1000 in 2% BSA. For anti-Calnexin, the dilution was used at 1:20,000 in 5% non-fat dry milk. The incubation was done overnight at 4°C.

#### Secondary antibody incubation

The dilution was used at 1:10,000 in PBS-T. Both HRP-conjugated (Jackson ImmunoResearch) and fluorescent conjugated secondary antibodies were used at same dilution. The incubation for secondary antibodies was done at RT for 1h. Washing was done by 1X-PBS.

#### Detection

The gels images were acquired using enhanced chemiluminescence (ECL) by treating the membranes in plastic wrap with Immobilon Forte Western HRP Substrate (Millipore. Cat. No.: WBLUF0100) and imaging on Amersham Imager 600 or iBright imaging system. The gel images for fluorescent conjugated antibodies were acquired on Odyssey Clx (LI-COR Biosciences). The gel images were processed on Image J.

### Reagent List and secondary antibodies

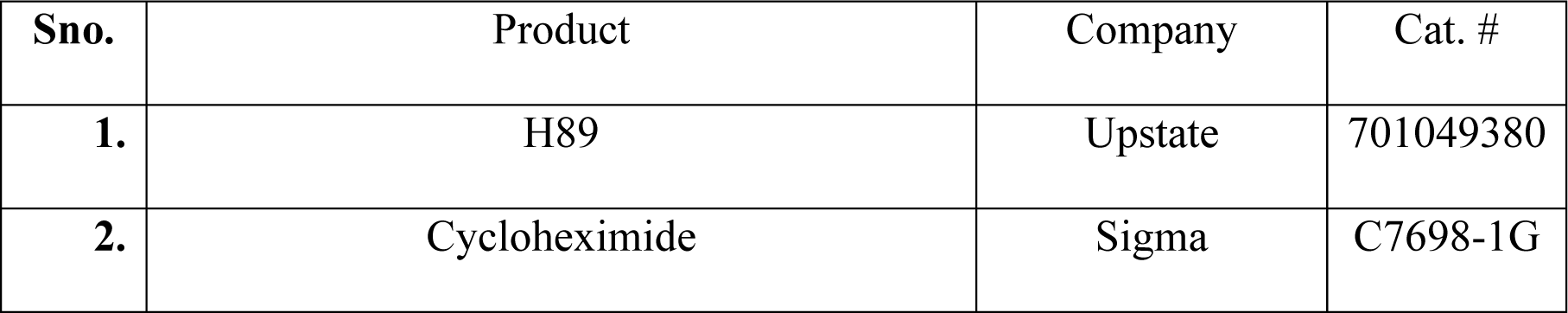
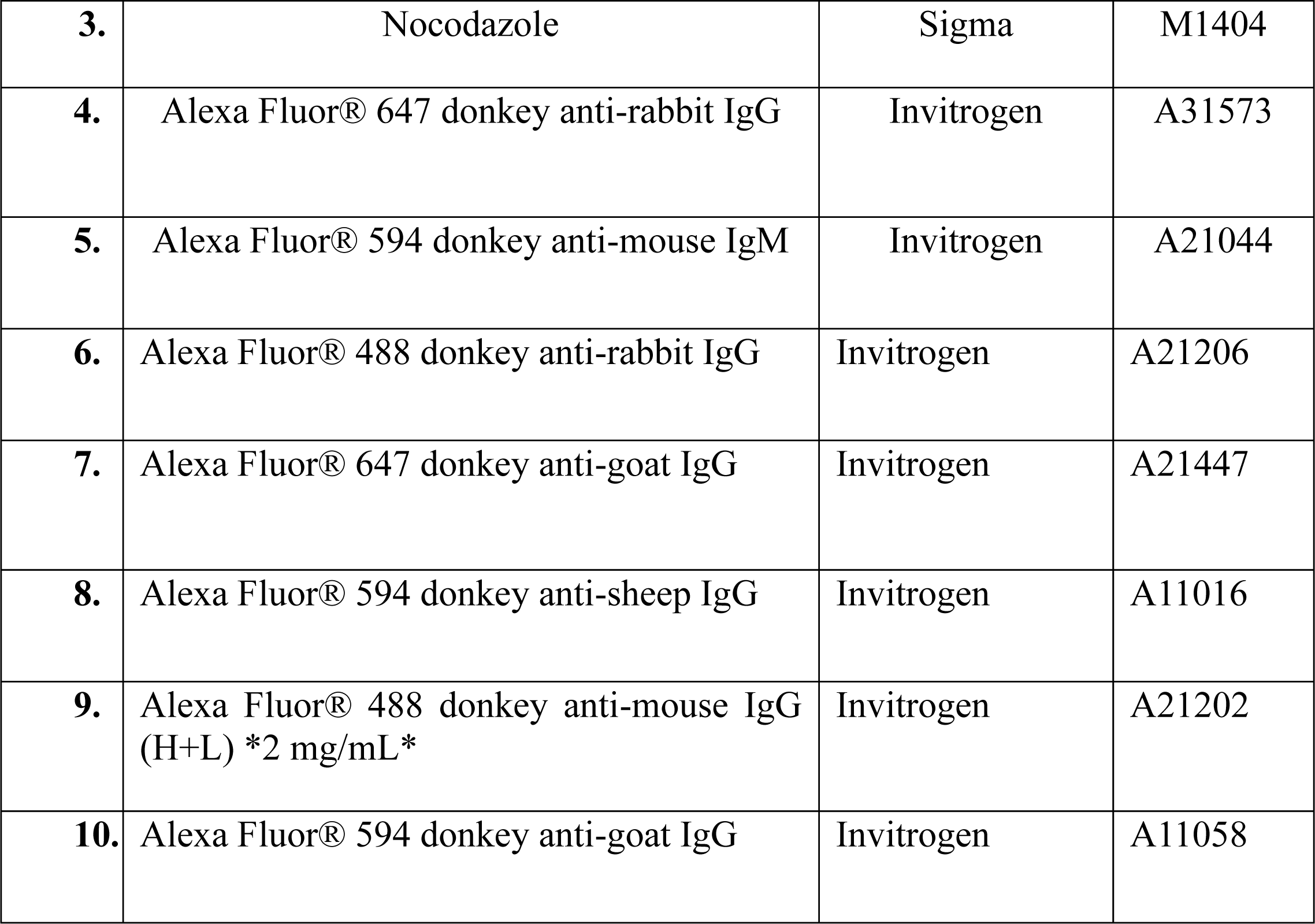

| Sno. | Product | Company | Cat. # |
| --- | --- | --- | --- |
| 1. | H89 | Upstate | 701049380 |
| 2. | Cycloheximide | Sigma | C7698-1G |
| **3.** | Nocodazole | Sigma | M1404 |
| **4.** | Alexa Fluor® 647 donkey anti-rabbit IgG | Invitrogen | A31573 |
| **5.** | Alexa Fluor® 594 donkey anti-mouse IgM | Invitrogen | A21044 |
| **6.** | Alexa Fluor® 488 donkey anti-rabbit IgG | Invitrogen | A21206 |
| **7.** | Alexa Fluor® 647 donkey anti-goat IgG | Invitrogen | A21447 |
| **8.** | Alexa Fluor® 594 donkey anti-sheep IgG | Invitrogen | A11016 |
| **9.** | Alexa Fluor® 488 donkey anti-mouse IgG (H+L) *2 mg/mL* | Invitrogen | A21202 |
| **10.** | Alexa Fluor® 594 donkey anti-goat IgG | Invitrogen | A11058 |

## Acknowledgements

We thank all the lab members of Malhotra laboratory for their valuable input, discussions, and suggestions during the project. We thank Dr. Davide Normanno, Dr. Felix Ruhnow, Dr. Nadia Halidi, Dr. Arrate Mallabiabarrena Ormaechea and Dr. Raquel García Olivas for their valuable advice on microscopy and optogenetics. SS thanks Roger Pons Lanau for his valuable input regarding data quantifications. We thank the CRG advanced light microscopy unit (ALMU), the Protein Technologies Unit, and the CRG/UPF Flow Cytometry Unit. We acknowledge the support of the Spanish Ministry of Science and Innovation to the EMBL partnership, the Centro de Excelencia Severo Ochoa and the CERCA Programme / Generalitat de Catalunya. We acknowledge financial support from the following sources: Ministerio de Ciencia e Innovación (Ramon y Cajal Felow, RYC-2016-20919) to OF, Ministerio de Economía y Competitividad: IJCI-2017-34751 to IR, SEV-2012-0208, BFU2013-44188-P, CSD2009-00016 to VM. VM also acknowledges the support by the European Research Council (ERC) synergy grant ERC-2020-SyG-951146. This publication is part of the Project TARTAFI (Ref. PDC2021-121870-I00), funded by MCIN/AEI/10.13039/501100011033 and the European Union “NextGenerationEU”/PRTR.

*Supplementary videos can be provided upon reasonable request.

## Supplementary Information

**Supplementary Figure 1:**
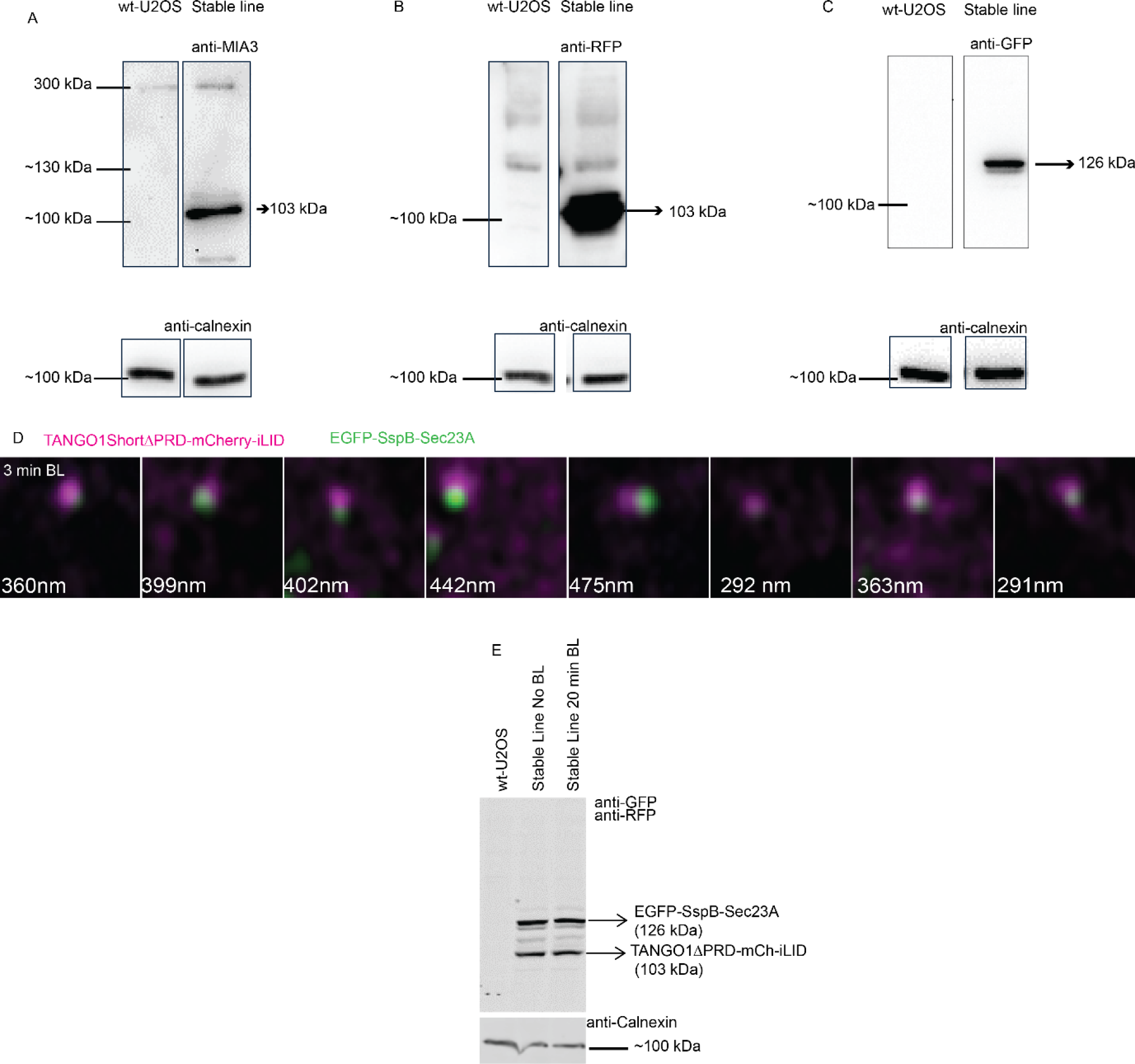
Generation of Stable cell lines in U2OS and their characterization: Lysates of wt-U2OS cells and stable line co-expressing TANGO1ShortΔPRD-mCh-iLID and EGFP-SspB-Sec23A were subjected to western blot analysis. Calnexin was used as control. **(A)** Top panel shows antibodies staining against MIA3 that binds to coiled coil domains of TANGO1 family of proteins. Band visible at 300 kDa corresponds to TANGO1 full length (visible in both wild-type U2OS and the stable line). Band at ∼103 kDa corresponds to TANGO1ShortΔPRD-mCh-iLID expressed in the stable line. Lower panel shows bands corresponding to calnexin as a control. (The gel lanes were rearranged for better clarity). **(B)** Top panel shows antibodies staining against RFP that binds to mCherry. Band visible at ∼103 kDa corresponds to TANGO1ShortΔPRD-mCh-iLID expressed in the stable line. **(C)** Top panel shows antibodies staining against GFP that binds to EGFP. Band visible at ∼126 kDa corresponds to EGFP-SspB-Sec23A expressed in the stable line. **(D)** Representative puncta marked by TANGO1ShortΔPRD-mCherry-iLID (magenta) and EGFP-SspB-Sec23A (green) formed after exposing stable lines (co-expressing the optogenetic constructs) with blue light for 3 min. **(E)** Lysates of control (wt-U2OS) and stable lines co-expressing TANGO1ShortΔPRD-mCh-iLID + EGFP-SspB-Sec23A exposed to blue light for 0 min and 20 min, and further subjected to western blot analysis. Calnexin was used as control. Band at 126 kDa and 103 kDa corresponds to EGFP-SspB-Sec23A and TANGO1ShortΔPRD-mCh-iLID respectively. No change or degradation is observed in the protein bands exposed to blue light.

**Supplementary Figure 2:**
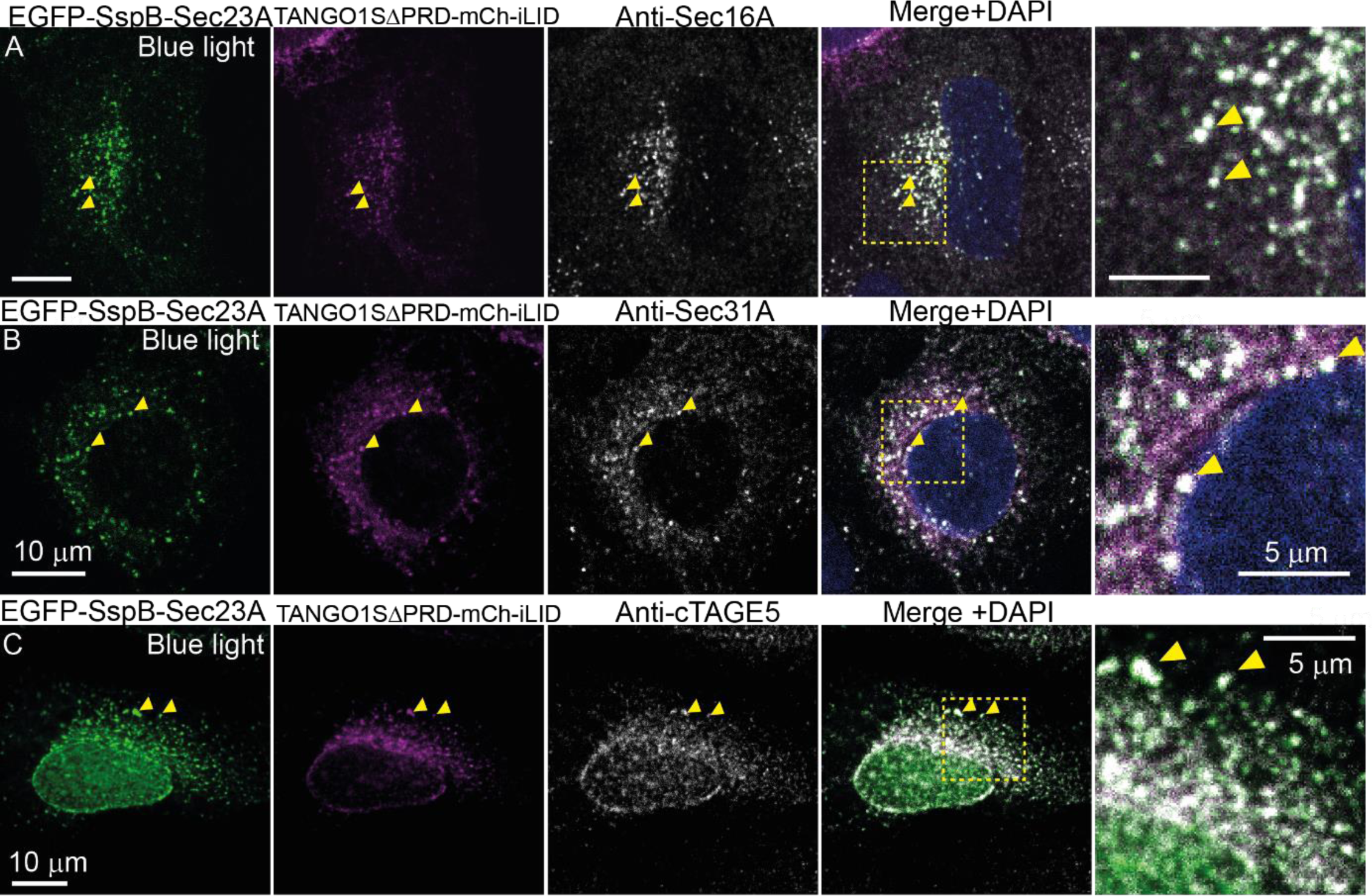
The puncta formed by TANGO1SΔPRD-mCh-iLID and EGFP-SspB-Sec23A are marked by ERES markers. Confocal images of U2OS cells transfected with TANGO1SΔPRD-mCh-iLID (magenta) and EGFP-SspB-Sec23A (green), exposed to blue light (1h), and fixed and stained for Sec16A, Sec31A and cTAGE5. Puncta formed by EGFP-SspB-Sec23A (green) and TANGO1SΔPRD-mCh-iLID (magenta) localized with (A) Sec16 (grey), (B) Sec31 (grey) and (C) cTAGE5 (grey), marked by yellow arrow heads.

**Supplementary Figure 3:**
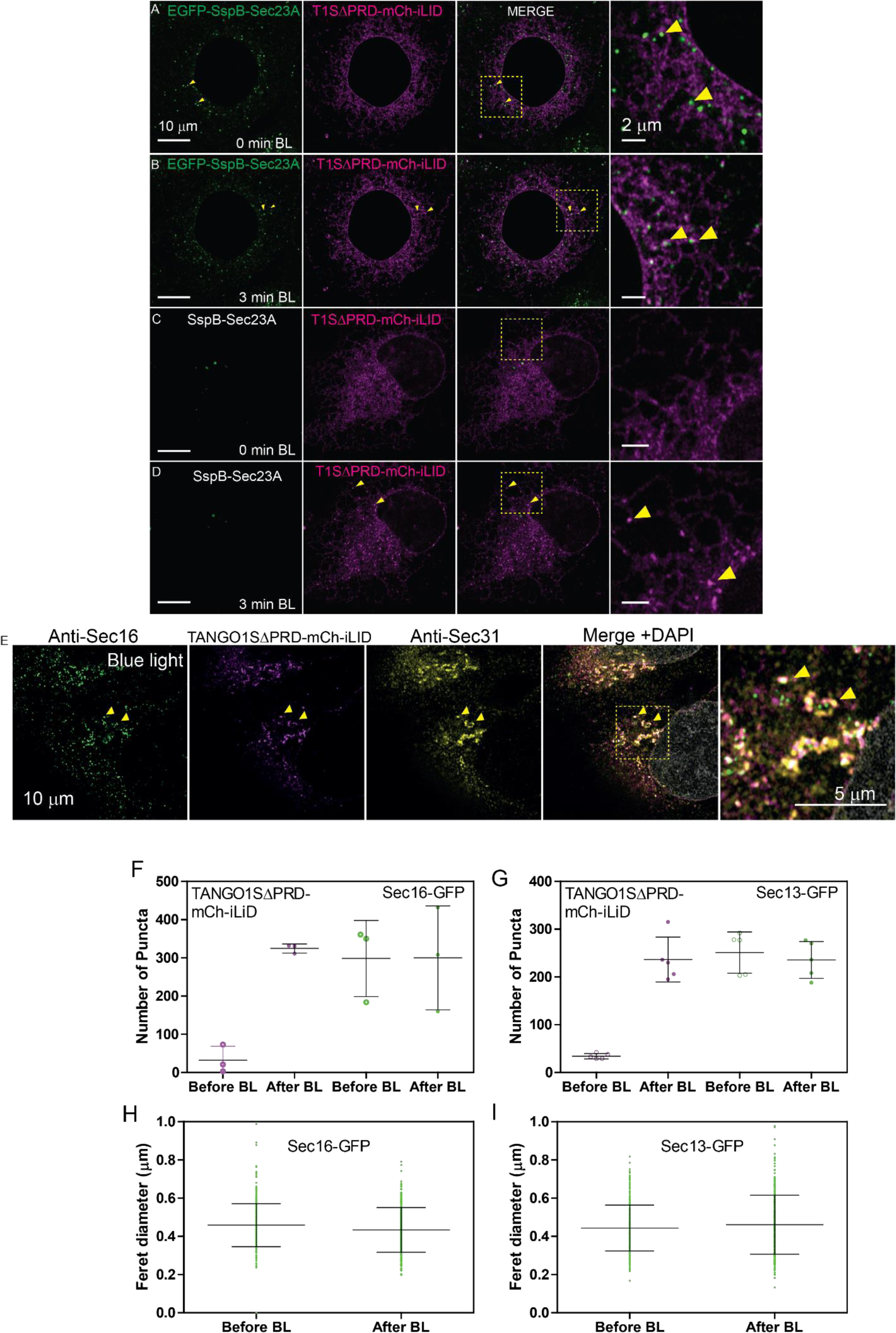
Generation of Stable cell lines in U2OS and their characterization: Snap shots of a time lapse of blue light irradiated stable line co-expressing TANGO1ShortΔPRD-mCherry-iLID (T1SΔPRD-mCh-iLID, magenta) + EGFP-SspB-Sec23A (green) at time **(A)** 0 min and **(B)** 3 min. Yellow arrowheads show the ERES marked by **(A)** EGFP-SspB-Sec23A, and **(B)** EGFP-SspB-Sec23A + TANGO1ShortΔPRD-mCherry-iLID (seen as white puncta). Snap shots of time lapse of another stable line (blue light irradiated) co-expressing TANGO1ShortΔPRD-mCherry-iLID (T1SΔPRD-mCh-iLID, magenta) + SspB-Sec23A at time **(C)** 0 min and **(D)** 3 min. Yellow arrowheads show the ERES marked by **(D)** TANGO1ShortΔPRD-mCherry-iLID bound to SspB-Sec23A (seen as magenta puncta). **(E)** Confocal image of stable line co-expressing TANGO1ShortΔPRD-mCherry-iLID (magenta) + SspB-Sec23A. Cells were exposed to blue light for 3 min, fixed and stained with antibodies against Sec16A (green) and Sec31A (yellow). ERESs marked with TANGO1ShortΔPRD-mCherry-iLID were enriched in Sec16A and Sec31A and are shown by yellow arrow heads. **(F)** A plot to compare the number of puncta formed by TANGO1SΔPRD-mCh-iLID (magenta) and Sec16L-GFP, before and after exposing with blue light (BL) for 3 min. The horizontal line corresponds to the mean, and error bar are shown. Data acquired from 3 cells in n=3 experiments. **(G)** A plot to compare the number of puncta formed by TANGO1SΔPRD-mCh-iLID (magenta) and Sec13-GFP, before and after exposing with blue light (BL) for 3 min. The horizontal line corresponds to the mean, and error bar are shown. Data acquired from 3 cells in n=3 experiments. **(H)** An aligned dot plot showing the ferret diameters of puncta formed by TANGO1SΔPRD-mCh-iLID (magenta) after 3 min blue light exposure and Sec16L-GFP (green) before and after 3 min blue light exposure. Feret sizes of 300 distinct puncta were used from at least n=3 data sets. **(I)** An aligned dot plot showing the ferret diameters of puncta formed by TANGO1SΔPRD-mCh-iLID (magenta) after 3 min blue light exposure and Sec13-GFP (green) before and after 3 min blue light exposure. Feret sizes of 300 distinct puncta were used from at least n=3 data sets.

**Supplementary Figure 4:**
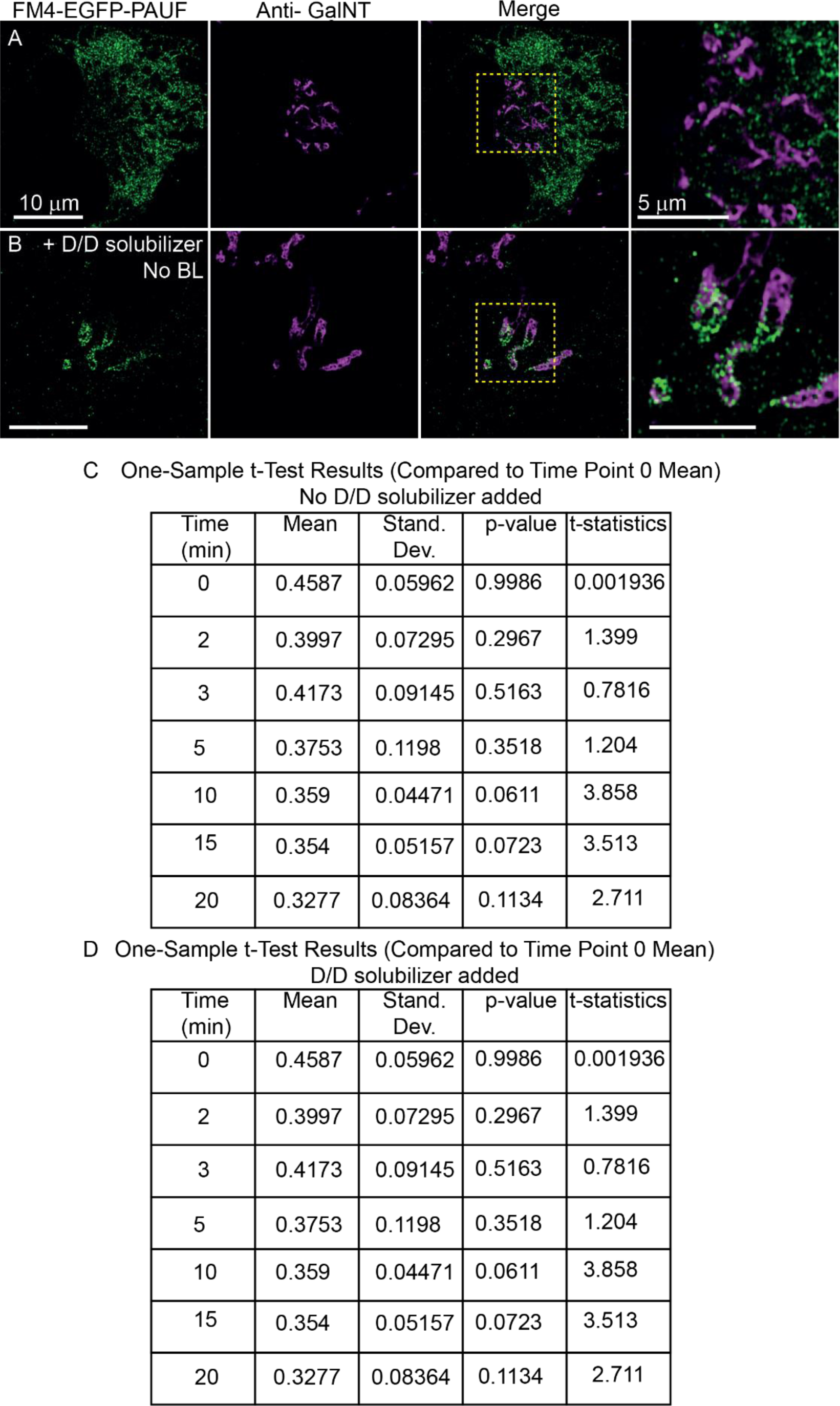
Fixed cells microscopy images of U2OS cells expressed with FM4-EGFP-PAUF (green) **(A)** untreated and **(B)** treated with D/D solubilizer for 20 min. Cells stained for antibody against GalNT (magenta). (Images acquired on Airyscan mode). **(A)** The FM4-EGFP-PAUF in aggregated state is trapped in the ER. The region marked by dotted yellow box shows Golgi region marked by GalNT and is devoid of FM4-EGFP-PAUF. **(B)** The addition of D/D solubilizer enables the dis-aggregation of the FM4-EGFP-PAUF, and now it is available for release from the ER. The FM4-EGFP-PAUF is observed accumulating at the Golgi cisternae membrane marked by GalNT after 20 min. (**C, D**) Table showing results of on-sample t-test for PCC of TANGO1-mCh-iLID and FM4-PAUF-EGFP for **(C)** No D/D solubilizer added and **(D)** D/D Solubilizer added.

**Supplementary Figure 5:**
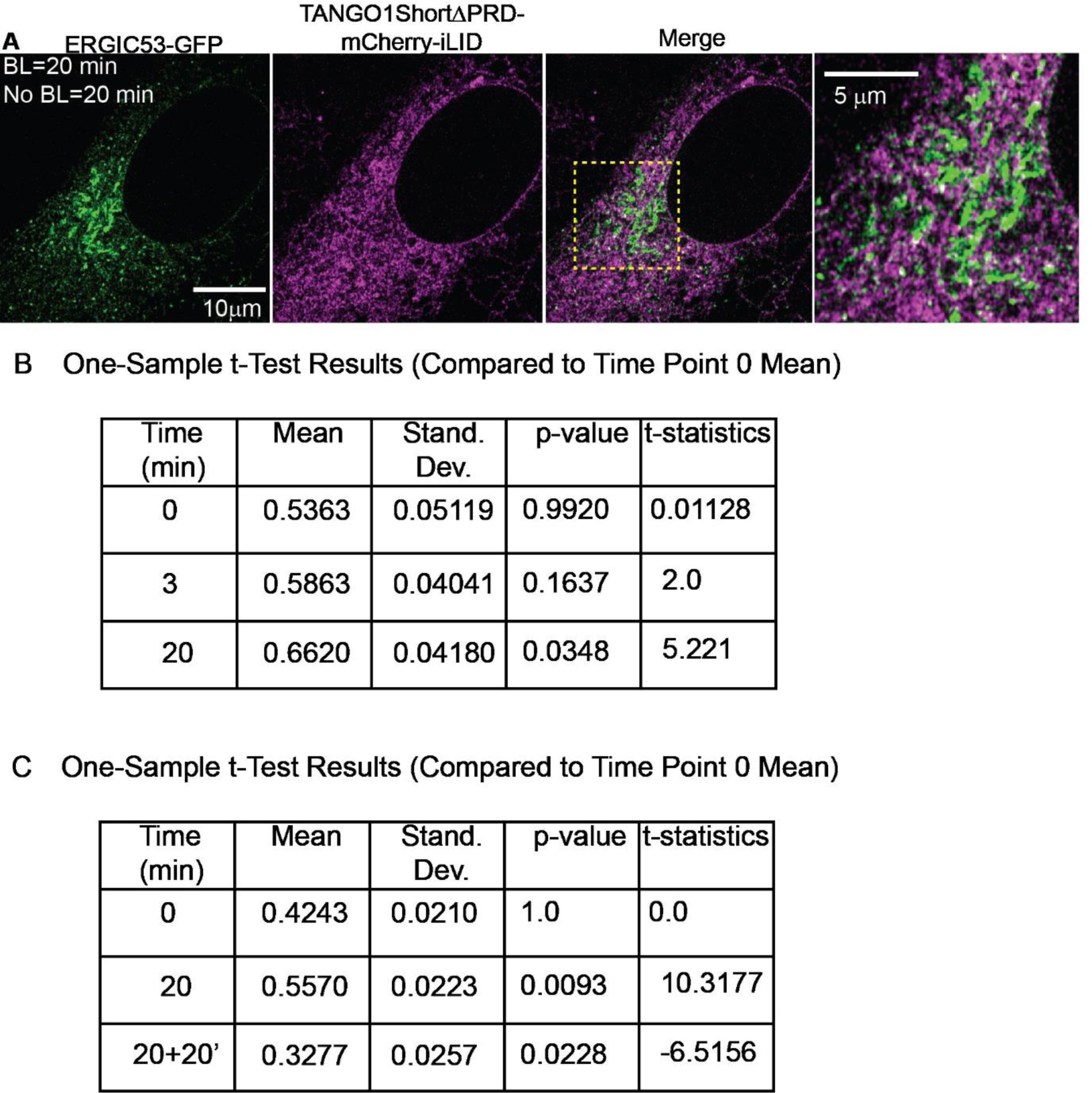
**(A)** Image of U2OS cell, stably co-expressing SspB-Sec23A and TANGO1SΔPRD-mCh-iLID, transfected with ERGIC53-GFP. Cells were exposed to blue light for 20 min and image was acquired after removing the blue light for another 20 min. The time lapse was acquired on Airyscan microscope showing ERGIC53-GFP (green) and TANGO1SΔPRD-mCh-iLID (magenta). (**B, C**) Table showing results of on-sample t-test for PCC of TANGO1-mch-iLID and ERGIC53-GFP in **(B)** absence and **(C)** presence of cycloheximide with exposure to blue light. Timepoint 20+20’ corresponds to 20 min blue light+ 20 min no blue light.

**Supplementary Figure 6:**
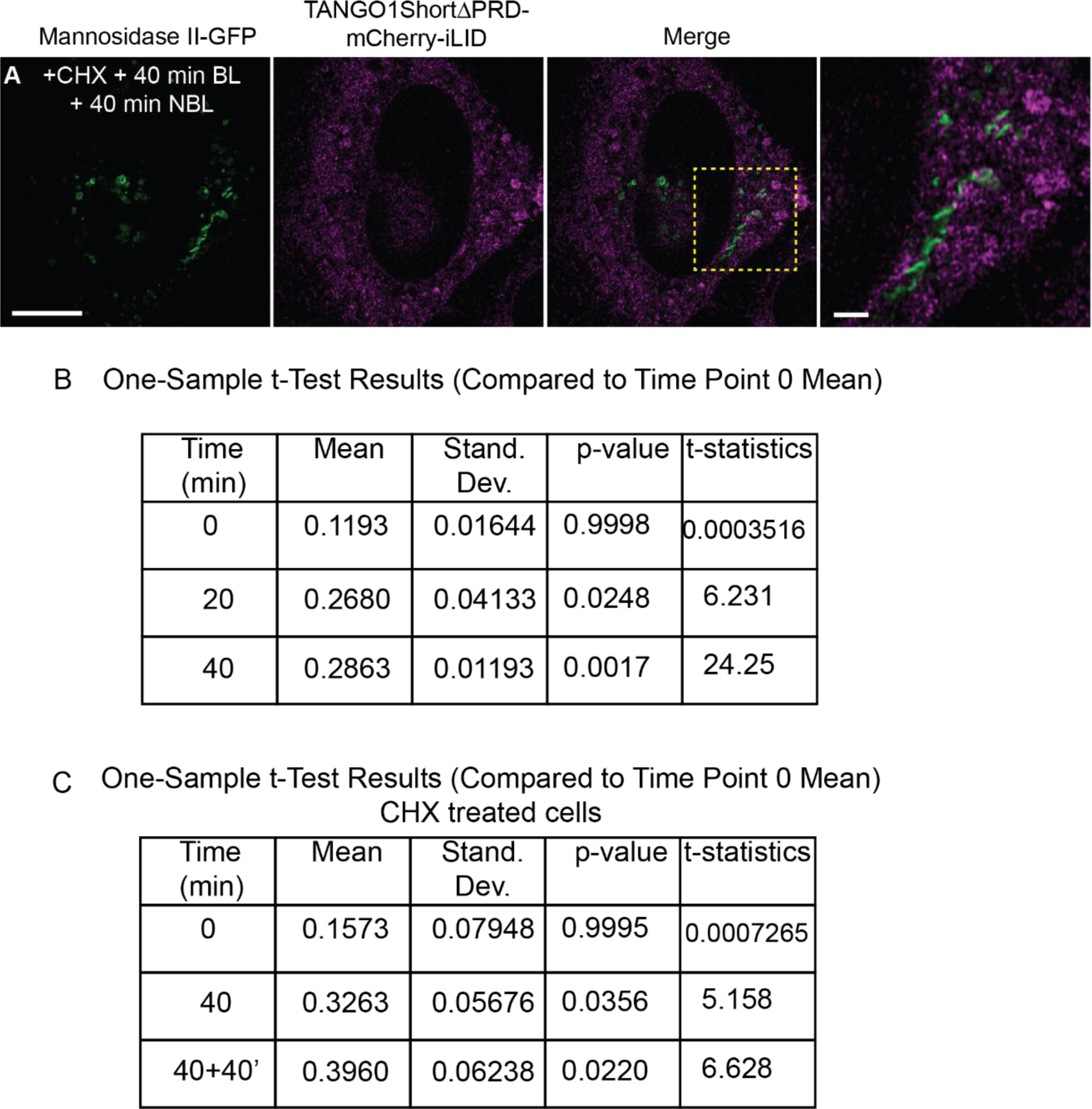
**(A)** Image of U2OS cells, stably co-expressing SspB-Sec23A and TANGO1SΔPRD-mCh-iLID, transfected with Mannosidase-II-GFP, and treated with cycloheximide (100 μM, for 30 min). Cells were exposed to blue light for 40 min and snapshot was acquired after removing the blue light for another 40 min. The image was acquired on Airyscan microscope showing Mannosidase-II-GFP (green) and TANGO1SΔPRD-mCh-iLID (magenta). **(B, C)** Tables showing the one sample t-tests results of mean of PCC of TANGO1ShortΔPRD-mCh-iLID and Mannosidase II-GFP, with blue light **(B)** in absence of cycloheximide and **(C)** in presence of cycloheximide. The results show mean, standard deviation, p-value, and t-statistics.

**Supplementary Figure 7:**
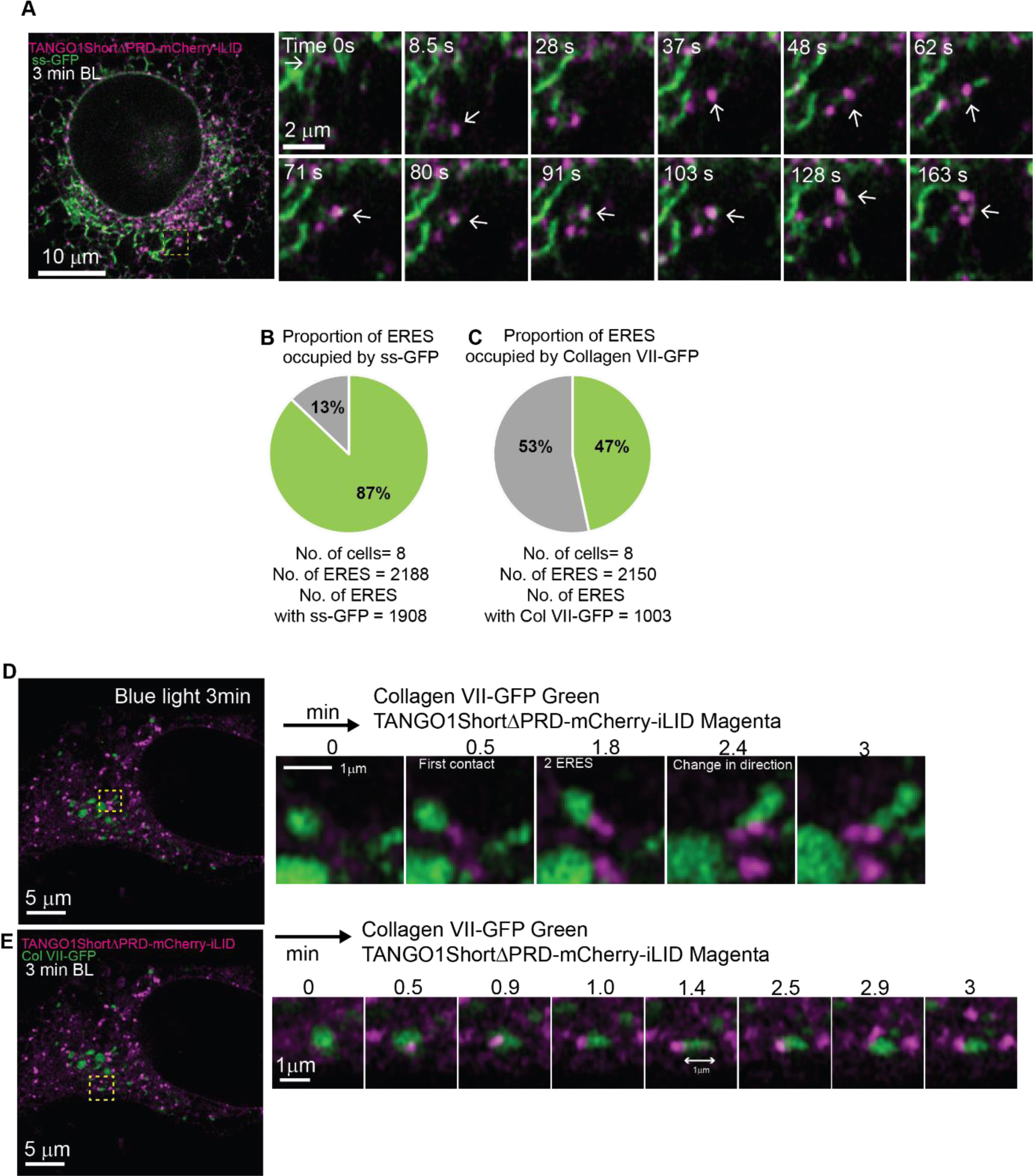
**(A)** Snapshots from time lapse of U2OS cells (Supplementary material video 8A), stably co-expressing SspB-Sec23A and TANGO1SΔPRD-mCh-iLID, transfected with ss-GFP. Interactions of ss-GFP with an ERES are shown with time in the zoom. **(B)** Pie-chart of proportion of vacant ERESs and ERESs occupied with ss-GFP. **(C)** Pie-chart of proportion of vacant ERESs and ERESs occupied with Collagen VII-GFP. **(D)** Snap shots of time-lapse (Supplementary material video 8C) of cell stably co-expressing TANGO1SΔPRD-mCh-iLID (magenta) and SspB-Sec23A transfected with Collagen VII-GFP (green), irradiated with blue light for 3 min. In zoom one representative ERES at various time intervals is shown. An ERES is observed in contact with collagen VII-GFP at 0.5 min. Another ERES appears at 1.8 min and the ERESs and collagen VII-GFP align on the plane. At 2.4 min the whole structure is observed to change its direction and collagen VII-GFP appears to extend in a tubular structure till end of time lapse. **(E)** Snap shots of time-lapse (Supplementary material video 8C) of cell stably co-expressing TANGO1SΔPRD-mCh-iLID (magenta) and SspB-Sec23A transfected with Collagen VII-GFP (green), irradiated with blue light for 3 min. In zoom a tubular collagen at an ERES at various time intervals is shown. Size of tubule of collagen VII -GFP is also shown.

**Supplementary Figure 8:**
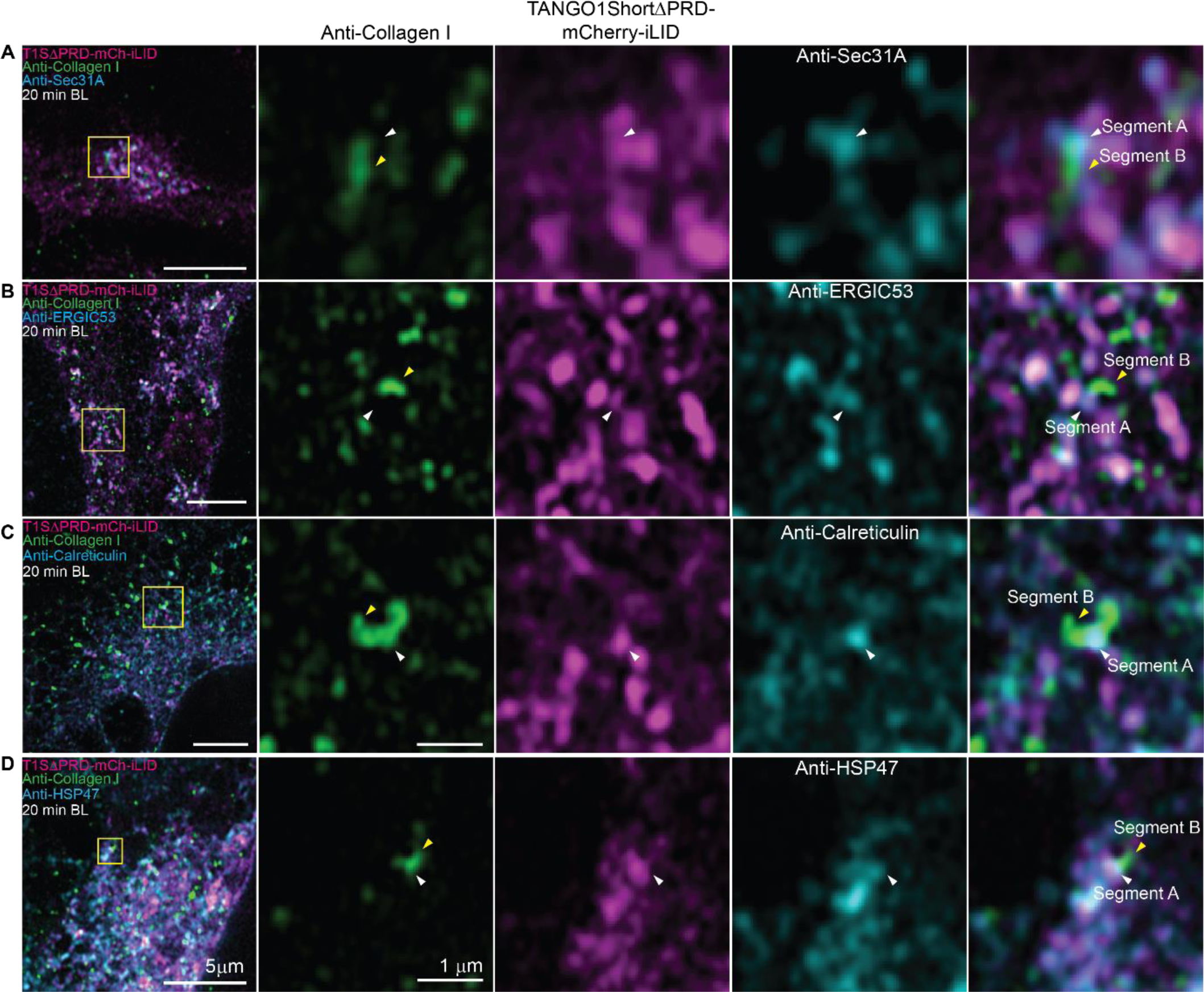
Microscopy image of U2OS cell co-expressing TANGO1SΔPRD-mCh-iLID (magenta) and SspB-Sec23A, irradiated with blue light for 20 min. Cells were fixed and stained with antibodies against endogenous collagen I and **(A)** Sec31A **(B)** ERGIC 53, **(C)** Calreticulin, and **(D)** HSP47. For TANGO1ΔPRDmCh-iLID, antibody against mCherry was used to enhance its signal. Endogenous Collagen I appear as tubules and puncta in the cell. The two segments of tubular collagen I at ERES are shown, ‘segment A’ that co-localizes with ERES (marked by white arrowhead) and ‘segment B’ that doesn’t colocalize with the ERES (marked by yellow arrowhead). **(A)** Zoomed image shows a tubular structure of collagen I, where segment A is localized with Sec31A (marked in white arrowhead) enriched with TANGO1SΔPRD-mCh-iLID. **(B)** Zoomed image shows a tubular structure of collagen I, where segment A is localized with ERGIC53 (marked in white arrowhead) enriched with TANGO1SΔPRD-mCh-iLID. **(C)** Zoomed image shows a tubular structure of collagen I, where segment A is localized with calreticulin (marked in white arrowhead) enriched with TANGO1SΔPRD-mCh-iLID, but the segment B is not localized with either. **(D)** Zoomed image shows a tubular structure of collagen I, where segment A is localized with HSP47 (marked in white arrowhead) enriched with TANGO1SΔPRD-mCh-iLID, but the segment B is not localized with either.

